# A Third Somatomotor Representation in the Human Cerebellum

**DOI:** 10.1101/2022.04.13.488247

**Authors:** Noam Saadon-Grosman, Peter A. Angeli, Lauren M. DiNicola, Randy L. Buckner

**Affiliations:** Department of Psychology, Center for Brain Science, Harvard University, Cambridge, MA 02138, USA; Athinoula A. Martinos Center for Biomedical Imaging, Massachusetts General Hospital, Charlestown, MA 02129, USA; Department of Psychiatry, Massachusetts General Hospital, Charlestown, MA 02129, USA

## Abstract

Seminal neurophysiological studies in the 1940s discovered two somatomotor maps in the cerebellum – an inverted anterior lobe map and an upright posterior lobe map. Both maps have been confirmed in the human using non-invasive neuroimaging with additional hints of a third map near to the cerebellar vermis. Here we sought direct evidence for the third somatomotor map by using intensive, repeated functional MRI (fMRI) scanning of individuals performing movements across multiple body parts (tongue, hands, glutes and feet). An initial discovery sample (N=4, 4 sessions per individual including 576 separate blocks of body movements) yielded evidence for the two established cerebellar somatomotor maps, as well as evidence for a third discontinuous foot representation near to the vermis. When the left versus right foot movements were directly contrasted, the third representation could be clearly distinguished from the second representation in multiple individuals. Functional connectivity from seed regions in the third somatomotor representation confirmed anatomically-specific connectivity with the cortex, paralleling the patterns observed for the two well-established maps. All results were prospectively replicated in an independent dataset with new individuals (N=4). These collective findings provide direct support for a third somatomotor map in the vermis of the cerebellum. We discuss the relations of this candidate third map to the broader topography of the cerebellum as well as its implications for understanding the specific organization of the human cerebellar vermis where distinct zones appear functionally specialized for somatomotor and visual domains.

## Introduction

A somatomotor topography was first described in the anterior lobe of the cerebellum by Lord Adrian (Adrian 1943). In cats and monkeys, Adrian recorded afferent discharges reaching the cerebellum following electrical stimulation to the cerebral cortex, tactile stimulation, movements of joints and stretching of muscles. He discovered that body parts are topographically organized in an inverted fashion beginning with the hindlimb represented in lobules III and HIII, the forelimb in lobules IV, V and HIV and HV, and the face in lobules VI and HVI at the posterior border of the anterior lobe. A year after Adrian’s seminal observations, Snider and Stowell (1944) confirmed the anterior lobe somatomotor topography and discovered a second discontinuous body map in the posterior lobe that was upright rather than inverted. They recorded cerebellar evoked potentials, also in cats and monkeys, following tactile stimulation. In the paramedian lobule (lobules HVII, HVIII), the hindlimb representation was found posteriorly, the face anteriorly and the forelimb in between (Snider and Stowell 1944; see also Snider and Eldred 1952).

A half century after their discovery, the anatomical basis of the two separate somatomotor maps was revealed using transneuronal viral tracing techniques. A barrier to measuring anatomical connectivity between the cerebral cortex and the cerebellum was that the pathways are polysynaptic (Evarts and Thach 1969; Kemp and Powell 1971; Strick 1985; Schmahmann and Pandya 1997) and did not yield to traditional monosynaptic anatomical tracing techniques. Using polysynaptic tracing techniques based on modified herpes and rabies viruses, Strick and colleagues (Middleton and Strick 1994; Kelly and Strick 2003) demonstrated that the hand region of the monkey’s cerebral motor cortex projects to and receives input from cerebellar lobules V-VI in the anterior lobe and HVIIb-HVIII in the posterior lobe, consistent with the two well-established somatomotor maps.

The two distinct somatomotor maps have been repeatedly observed in humans using non-invasive neuroimaging techniques evoked by movements (Nitschke et al. 1996; Rijntjes et al. 1999; Grodd et al. 2001; Buckner et al. 2011; Diedrichsen and Zotow 2015; Guell et al. 2018a) and through tactile stimulation (Fox et al. 1985, Bushara et al. 2001; Takanashi et al. 2003). Functional connectivity between cerebral motor regions and the cerebellum in task-free data also reveals the two maps (Buckner et al. 2011; Guell et al. 2018b; Xue et al. 2021). Thus, multiple approaches have converged to suggest that the cerebellum possess two distinct somatomotor body maps.

Beyond these two established somatomotor body maps, we have also postulated the existence of a third, smaller body map in the most posterior extent of the cerebellum near to the vermis (Buckner et al. 2011). There are multiple motivations for this hypothesis. In comprehensive mapping efforts that have explored both motor and nonmotor regions of the cerebellum, an orderly progression of zones is found along the anterior-to-posterior axis of the cerebellum variably described as a sequence of networks or a gradient (Buckner et al. 2011; Buckner 2013; Guell et al. 2018b; Xue et al. 2021). The progression begins with the inverted somatmotor map in the anterior lobe of the cerebellum and then progresses through multiple sensory-motor and higher cognitive networks ending at the Crus I / II border. The progression of networks inverts at the Crus I / II border and proceeds in mirror-reversed order through the posterior lobe of the cerebellum ending on the well-established upright somatomotor map. While these two gradients alone capture many features of the observed organization of the cerebellum, they fail to explain a key feature.

Specifically, cerebral networks associated with higher-order functions display representations in the posterior lobe of the cerebellum near lobule IX, which would be expected if there was a third gradient, including somatomotor representation. In particular, the network referred to in the human literature as the default network possesses a robust representation near lobule IX (e.g., Habas et al. 2009; Krienen and Buckner 2009; Buckner et al. 2011; Xue et al. 2021). Tasks targeting social inferences, which increase response in the default network, also reveal a response in lobule IX (King et al. 2019; see also Diedrichsen and Zotow 2015; Guell et al. 2018a). Furthermore, in their seminal work, Kelly and Strick (2003) noted that Crus I / II receive prefrontal projections but also described a smaller group of neurons in lobules IX / X. The labelled neurons fall within the region of the suggested tertiary association map in the posterior end of the cerebellum (see Guell et al. 2018a Figure 1). One possibility is that a third map represents cognitive networks but not motor networks, or what Guell et al. (2018a) refer to as the “double motor/triple nonmotor representation hypothesis.” The alternative possibility is that the cerebellum possesses a third somatomotor map that is small and challenging to identify.

**Fig. 1.**
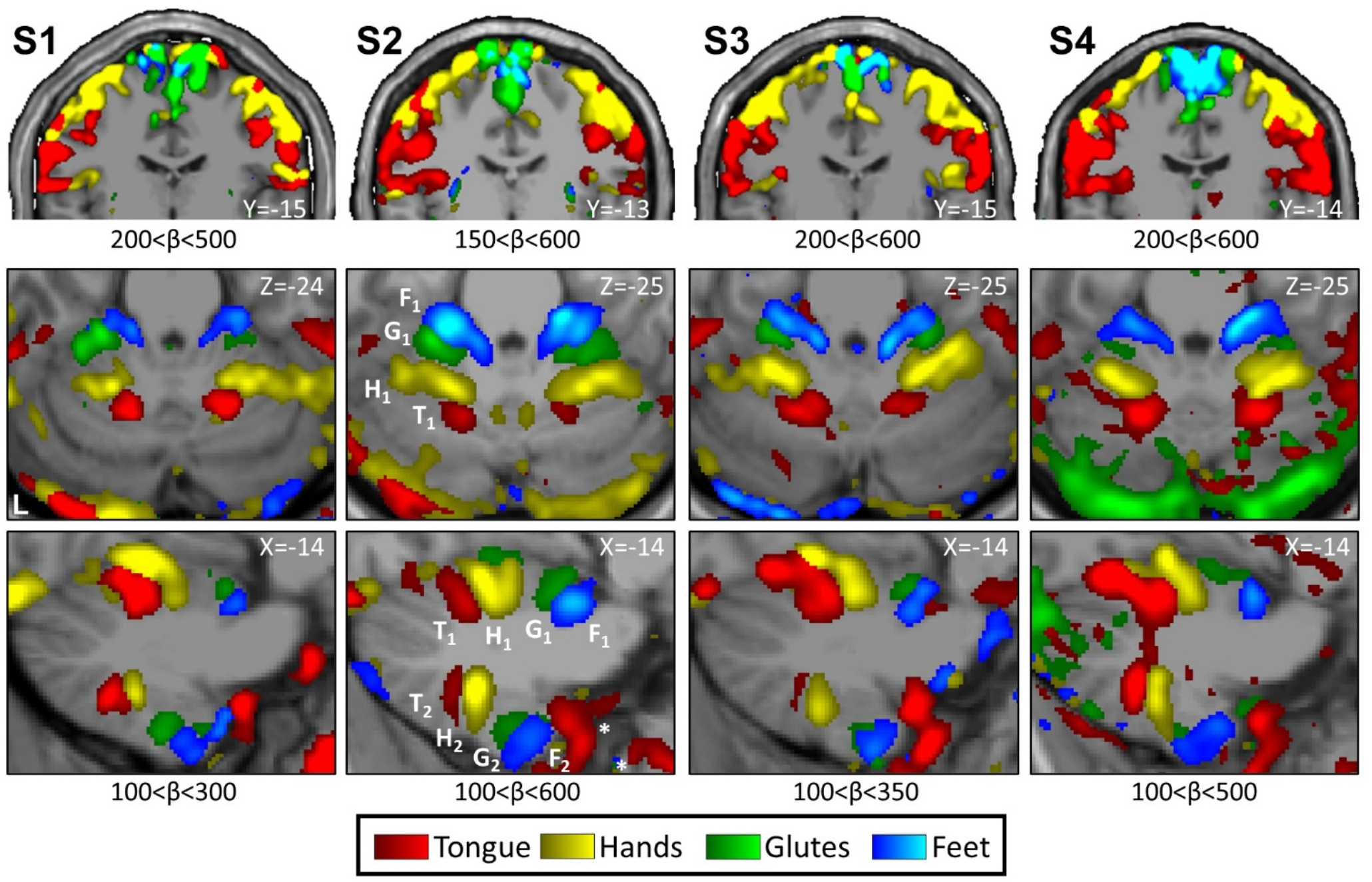
Somatomotor topography in the cerebral cortex and cerebellum is evident in individual participants. Winner-take-all maps of active movements are displayed for 4 body parts along the anterior-posterior body axis: tongue (red), right and left hands (yellow), glutes (green) and right and left feet (blue). Each column displays a separate participant (S1-S4). Beta values are thresholded to best capture topography, separately for the cerebral cortex and cerebellum. In each participant a clear body topography is evident by the order-tongue(T)-hand(H)-glutes(G)-foot(F) in the primary motor cortex, M1 (lateral to medial, top row), cerebellar anterior lobe (middle and bottom rows) and in the cerebellar posterior lobe (bottom row). The primary cerebellar representation (T1-H1-G1-F1) is inverted to the secondary representation as labelled in S2 (T2-H2-G2-F2). Note the tongue assignment anterior to the secondary representation (in all participants, marked with an asterisk in S2) is outside the cerebellum and likely results from a motion artifact. Coordinates indicate the section level in the space of the MNI152 atlas. The color bars represent beta values. L indicates left.

Hints of a third somatomotor representation exist in the literature. In an early electrophysiological study, Dow and Anderson (1942) documented responses to tactile stimulation in the pyramis (part of the posterior vermis) of rats. With transneuronal viral tracing techniques, the cerebellar vermis was shown to be a target of projections from motor areas in the cerebral cortex of monkeys (Coffman et al. 2011). In humans, one of the first studies to use fMRI to map the cerebellum noted that there might be a third somatomotor representation in the vermis (Rijntjes et al. 1999). However, this suggestion was made with caution due to the small sample size and mixed results. In our prior well-powered, group-averaged study, a small medial region near to the vermis of the cerebellum was linked to cerebral somatomotor networks (see the transverse section at Z = −38 in Figure 8 of Buckner et al. 2011). But the representation in the vermis could not be disambiguated from the adjacent zones of the cerebellum attributed to the established second somatomotor map. Similarly, Guell et al. (2018a) noted a candidate motor representation of the foot near the vermis but with sufficient location uncertainty to downplay the result.

A definitive demonstration of a third somatomotor map, if one exists, faces multiple challenges. First, the hypothesized map is expected to be found in a small area within the already small cerebellum. Second, the inferior portions of the cerebellum near the vermis have low signal-to-noise ratio (SNR) in fMRI compared to the cerebral cortex.^1^ Finally, a third somatomotor map is expected to be found in close spatial proximity to the second map, making it difficult to dissociate between the two posterior lobe representations.

In the present work we took a number of steps to overcome these challenges. First, we acquired a substantial amount of data within individual participants during task-based motor movements to boost SNR. Each participant was scanned on four days with six motor runs per day, for a total of 576 separate blocks of motor movements (96 blocks for each of six separate motor conditions). Second, to avoid between-individual spatial blurring, each participant was analyzed separately (Marek et al. 2018; Xue et al. 2021). Third, we employed an experimental paradigm that targeted movement across the body parts, including the hands and feet on both sides. Given the orientation of the second somatomotor map, one but not both of the hand and foot representations should juxtapose one another, allowing the other body part to demonstrate separation of the second and hypothesized third maps. As the results will reveal, we were able to identify a clear third map representation in most of the individuals studied, with movements of the feet being the critical condition to differentiate the second and third somatomotor maps.

## Methods

### Participants

Eight healthy adults, aged 19-25 [mean= 22.4 yr (SD= 2.6), 2 men, 7 right-handed], were recruited from the Boston area. Four participants contributed data to the initial Discovery sample, and 4 participants contributed data to the Replication sample. All participants were screened to exclude a history of neurological and psychiatric illness or ongoing use of psychoactive medications. Participants provided written informed consent through a protocol approved by the Institutional Review Board of Harvard University.

### MRI data acquisition

Scanning was conducted at the Harvard Center for Brain Science using a 3T Siemens Magnetom Prisma-fit MRI scanner and a 64-channel phased-array head-neck coil (Siemens Healthcare, Erlangen, Germany). Inflatable padding provided comfort and helped immobilize the head. Participants viewed a rear-projected display through a mirror attached to the head coil. Before each session, the screen presentation was adjusted so that the center point could be comfortably viewed without any eye strain. Participants’ eyes were monitored and video-recorded using the Eyelink 1000 Core Plus with Long-Range Mount (SR Research, Ottawa, Ontario, Canada), and alertness was scored during each functional run.

All participants were scanned across 4 MRI sessions on separate non-consecutive days. Six task-based runs were acquired each day where participants made active movements (motor runs) as well as 2 runs where participants fixated a centrally presented black crosshair on a light grey background (fixation runs). In total, each participant had 24 motor and 8 fixation runs. 3 participants (S5-S7) were scanned in an additional session not used here.

Functional data employed a multiband gradient-echo echo-planar pulse sequence sensitive to blood oxygenation level-dependent (BOLD) contrast (Moeller et al. 2010; Feinberg et al. 2010; Setsompop et al. 2012; Xu et al. 2013), generously provided by the Center for Magnetic Resonance Research (CMRR) at the University of Minnesota. The 4 Discovery sample participants (S1-S4) were scanned with two different spatial resolutions: 1.8mm (2 sessions) and 2.4mm (2 sessions) isotropic voxels (order of sessions was balanced across participants). All sessions of the 4 Replication sample participants (S5-S8) were scanned with 2.4mm isotropic voxels. Acquisition parameters for 1.8mm resolution: TR = 2000ms, TE = 30ms, flip-angle = 80°, matrix 122 × 122 × 87 (FOV-221×221), multislice 3× acceleration. 211 volumes were acquired for each fixation run and 214 volumes for each motor run. Acquisition parameters for 2.4mm resolution: TR = 1000ms, TE = 33ms, flip-angle = 64°, matrix 92 × 92 × 65 (FOV-221×221), multislice 5× acceleration. 422 volumes were acquired for each fixation run and 428 volumes for each motor run. The two resolutions in the Discovery sample were acquired to explore the possibility of higher (1.8mm) resolution disambiguating functional details better than lower (2.4mm) resolution. For the present purposes, the two resolutions were combined and smoothed with a uniform kernel. The first 2 sessions of S5 and the first session of S6 were acquired in a different FOV (211×211); therefore, the matrix for both BOLD runs and field maps was: 88 × 88 × 65 and BOLD TE = 32.6ms. The change in FOV, which occurred to accommodate larger heads, did not affect the quality of registration or impact the analyses in any way we could detect.

A T1-weighted structural image was obtained in each session using a rapid multiecho magnetization-prepared rapid gradient echo (MPRAGE) three-dimensional sequence (van der Kouwe et al. 2008): TR = 2200ms, TE = 1.57, 3.39, 5.21, 7.03ms, TI = 1100ms, flip-angle = 7°, voxel size 1.2mm, matrix 192 × 192 × 176, in-plane generalized auto-calibrating partial parallel acquisition (GRAPPA) 4× acceleration. In each session two dual gradient-echo B0 field maps were also acquired to correct for susceptibility-induced gradient inhomogeneities with slice spatial resolution matched to the BOLD sequence (1.8mm or 2.4mm). Other field map parameters included: 1.8mm resolution – TE = 4.73ms, 7.19ms, TR = 806ms, flip angle = 55°; 2.4mm resolution – TE = 4.45ms, 6.91ms, TR = 295ms, flip angle = 55°.

Exclusion criteria included a maximum absolute motion of no more than 1.8 mm. 1 motor run was excluded for S3, 2 motor runs for S4, 1 motor and 2 fixation runs for S6 and 1 fixation run for S8. In 3 motor runs of S3, an artifact in the form of a peak translation (in the z axis and to a lesser extent in the y axis) was spotted in approximately the same time point across these 3 runs. Maximum absolute motion did not exceed the set threshold for these runs (0.72, 1.01, 0.85mm), but we conservatively excluded them from analysis. Runs were excluded based on BOLD data quality prior to examination of task response patterns to avoid bias.

### Motor task

Participants performed a blocked-task paradigm consisting of 6 types of movements that targeted four separate parts along the anterior-posterior body axis: (1) *Tongue*: participants moved their tongue from right to left, touching their upper premolar teeth, (2,3) *Right and Left Hands*: participants moved their fingers alternating between tapping the thumb with the index finger and the thumb with the middle finger, (4) *Glutes*: participants contracted and relaxed their gluteal muscles, and (5,6) *Right and Left Feet*: participants alternated dorsiflexion and plantarflexion of their toes.

Each movement was performed repeatedly in 10s active movement blocks. Prior to a movement block, a visual cue of a drawn body part with a text label was presented for 2s, informing the participant to initiate one of the six movement types. Then, the fixation crosshair was presented with a black circle surround that repeatedly flashed on and off to pace the movements (1s on then 1s off). The onset of the black circle cued movement of the tongue to the right, thumb to index, glutes contraction and toes plantarflexion. The black circle turning off cued movement of the tongue to the left, thumb to middle, glutes relaxation and toes dorsiflexion. The word ‘End’ was presented for 1s at the end of each movement block to instruct the participant to stop, and there was a 1s fixation gap before the onset of the next movement cue.

Twenty-four movement blocks (4 per body part) occurred within each run, with 16s blocks of passive fixation providing a break after each set of 6 movement blocks. All motor runs began and ended with fixation yielding 5 fixation blocks per run. Six separate runs with distinct orders of movement conditions were performed on each day. Counterbalancing of the movement conditions across the 6 runs on each day ensured that each condition appeared exactly once in each of the run positions.

Participants extensively practiced the intended movements before the first scanning session. First, participants practiced remotely during a consent and training session, after watching a video demonstrating how to execute the movements. Emphasis was placed on how to localize each movement in a subtle way to avoid head motion. Next, when participants arrived for their first session, they practiced again while lying on the scanner bed outside of the bore and then a final time inside the bore. To further reduce extraneous motion, participants’ legs were supported in a semiflexed position using an ergonomic knee-to-ankle cushion.

### Data processing

A custom analysis pipeline for individualized data processing was used as described in detail in Braga et al. (2019). Briefly, the pipeline combines tools from FreeSurfer (Fischl 2012), FSL (Jenkinson et al. 2012) and AFNI (Cox 1996) to align data within an individual across runs and sessions to a high-resolution output target (1mm isotropic) using a single interpolation to minimize spatial blurring. Five different registration matrices were combined for the single interpolation: (1) a motion correction matrix for each volume to the run’s middle volume (linear registration, 12 degrees of freedom (DOF); MCFLIRT, FSL), (2) a matrix for field-map-unwarping the run’s middle volume (FUGUE, FSL), (3) a matrix for registering the field-map-unwarped middle volume to a mean BOLD template (12 DOF; FLIRT, FSL), (4) a matrix for registering the mean BOLD template to the participant’s native space T1 image (6 DOF; using boundary-based registration, FSL) and (5) a matrix for registering the native space T1 to the MNI152 1mm atlas (nonlinear registration; FNIRT, FSL). The mean BOLD template was created by taking the mean of all field-map-unwarped middle volumes after registration to an upsampled (1.2mm), unwarped mid-volume template (temporary target, selected from a low motion run acquired close to a field map). The native space template was one of the participant’s T1 structural images, upsampled to 1mm isotropic resolution. Given that multiple structural images were available for each individual, a single reference image was chosen that possessed a robust estimate of the pial and white-matter boundaries (as constructed by FreeSurfer recon-all).

Data were checked for registration errors. In cases of sub-optimal registration, a different mid-volume temporary template was chosen, or a different/additional field map was used for unwarping (since two field maps were acquired in each session). Before the calculation of these matrices, for each BOLD run, the first 12 volumes were discarded for T1 equilibration (6 volumes in 1.8mm resolution). For the participants (S1-S4) that had two spatial resolutions (two sessions with 1.8mm isotropic voxels and two sessions with 2.4mm isotropic voxels), each resolution was processed separately and then combined. All functional data were smoothed in volume with a 3mm full width at half maximum (FWHM) kernel.

Data were analyzed in the standard space of the MNI152 atlas within each participant individually. Thus, all idiosyncratic details within the individual were preserved. The use of the atlas space allowed the separate participants to be examined in a spatially registered framework and also allowed their data to be projected to a flat representation of the cerebellar cortex to aide visualization of patterns.

### Visualization on the cerebellar surface

To visualize the spatial extent of maps in the cerebellum, the data were projected onto a flat representation of the cerebellar surface (similar to our prior work in Xue et al. 2021). This representation was created by Diedrichsen and Zotow (2015) and utilized within the spatially unbiased infratentorial template (SUIT) toolbox (http://www.diedrichsenlab.org/imaging/suit_fMRI.htm). The projection method uses an approximate surface of the grey and white matter defined on the SUIT template (here specifically, MNI152, normalized by FSL). The vertices on the two surfaces come in pairs. When projecting, the algorithm samples the data along the line connecting the two vertices and then takes the mean of these numbers to determine the value of the vertex.

The surface representation was used for display purposes only. The surface is not unique to each individual. Nonetheless, the surface representation is able to visualize the individual participant’s topographic details in a uniform, comprehensive display framework.

### Motor task analysis

Motor run data in MNI152 atlas space were analyzed within each individual separately. A high-pass filter with a 100-s (0.01-Hz) cut-off was applied to remove low-frequency drifts within each run. Run specific general linear model (GLM, using FEAT; FSL) analyses were performed, including the six movement conditions as regressors of interest. The onsets of the six movement blocks were modeled 1s before the initiation of movement (1s after the appearance of the visual cue). The duration of each event block was set to 10s. Events were modeled with a canonical hemodynamic response function (double-Gamma) along with its temporal derivative. GLM analyses resulted in voxel-level beta values for each movement compared to the run means.

### Somatomotor topography

Initial analyses focused on estimating the somatomotor map across the four body parts in the cerebral cortex and separately in the cerebellum. A contrast map was computed for each movement by subtracting the mean of all other movements’ beta values. This was done in each motor run separately and then averaged across runs (including averaging across resolutions in S1-S4). To visualize somatomotor topography, a winner-take-all map was generated. Each voxel was assigned the highest beta value across all movement contrasts (one versus all the others) and was colored with the color of the corresponding movement: tongue-red, right and left hand-yellow, glutes-green, and right and left foot-blue. In this visualization, right and left representations were not distinguished. Maps were visualized separately for each individual in the volume and also projected onto the SUIT flatmap template of the cerebellum. Maps were thresholded within individuals to best capture somatomotor topography, separately for the cerebrum and for the cerebellum.

### Focus on the hand and foot representations

A critical aspect of the design is the ability to contrast left and right movements separately for the hands and feet. This enables direct contrast of the left and right movements within each body part, allowing a clean estimate of the representation of the body part because non-specific noise and other features of the data are largely nulled (paralleling the strategy we have repeatedly used in group-averaged functional connectivity data; Krienen and Buckner 2009; Buckner et al. 2011). Contrast maps were averaged across runs (and across resolutions in S1-S4). Contrast maps were projected onto a flatmap template of the cerebellar surface and thresholded within individuals.

### Seed based functional connectivity analysis

To assess representations’ specificity, we conducted a seed based functional connectivity analysis. This analysis should not be considered independent as we utilized all available data, including the motor task runs used to make the contrast maps. Specifically, in addition to the 24 motor runs (across 4 sessions) of each individual, we used the 8 fixation runs (2 per session), resulting in 32 possible runs. After quality control exclusions, S3 had 28 runs (193.3min), S4 had 30 runs (207.2min), S6 had 29 runs (200.5min) and S8 had 31 runs (214.2min). All other participants utilized all 32 runs (221.1min). Note that unlike purely task-free data, the present data included substantial modulation due to active task demands.

Pre-processing included all steps as used for the prior task-based analyses but also regression of nuisance variables (3dTproject, AFNI) as is typical in functional connectivity analysis (whole brain, ventricular, and deep cerebral white matter mean signals, as well as six motion parameters, and their temporal derivatives). The residual BOLD data were then band-pass filtered at 0.01–0.1 Hz and smoothed with a 3mm FWHM kernel paralleling the task-based analysis.

Seed regions were defined in the right and left primary, secondary and tertiary somatomotor cerebellar representations. For each seed region, an MNI152 coordinate was selected in the volume to best capture the center of the representation in the right versus left hand and foot contrast maps. In one individual, S8, we selected right and left foot tertiary coordinates to be similar to the ones of other individuals given the lack of detectable representations.

Seed regions were defined with a 2mm-radius centered at the selected MNI152 coordinate (33 voxels). For each run, the Pearson’s correlation was computed between the averaged time course of the seed region and that of all other voxels. The functional connectivity values were averaged across all runs (and resolutions for S1-S4) after being Fisher’s r-to-z transformed.

### Discovery and replication

A key aspect of our methods was prospective replication. Data from the initial four participants were fully analyzed and graphed (Discovery sample, S1-S4) before any analysis was attempted on the second, independent group of participants (Replication sample, S5-S8).

## Results

### Somatomotor topography is detected within the primary motor cortex

The body map in the primary motor cortex (M1, precentral gyrus) is well established. The anterior (head) to posterior (foot) axis are topographically organized as one moves from the lateral extent of the motor strip to the midline (Penfield and Boldrey 1937). Data from each individual revealed the full M1 body topography: tongue-hand-glutes-foot, lateral to medial as expected (Fig. 1, top). Although not common in the literature, contraction and relaxation of the gluteal muscles was included to evoke a representation of the trunk. As predicted by the body topography, representation of the glutes fell between the hand and the foot. The observed M1 motor topography validates the experimental paradigm and the processing methods applied.

### Somatomotor topography is detected in both of the established maps within the cerebellum

The two well-established cerebellar somatomotor maps were apparent in all participants (Fig. 1, middle and bottom). The ordering of body parts was inverted between the two maps. In the anterior lobe the foot was anterior to the tongue and in the posterior lobe the tongue was anterior to the foot (Fig. 1 bottom, highlighted in S2). In both maps, the glutes representation falls between the hand and foot filling in a gap often observed in motor body maps (e.g., Xue et al. 2021, their Fig. 3), although more apparent in the primary map than the secondary, and more clean for some participants (S1 and S2) than others (S3 and S4). The current data thus reveal strong and reliable evidence for the anterior lobe and posterior lobe somatomotor maps.

One other aspect of the results (Fig. 1 bottom) is that the tongue movements induced noise centered outside of the cerebellum (but extending into the cerebellum). Given the sensitivity of fMRI to motion artifact, it is not surprising that movements within the field of view produce artifact. Fortunately, the predicted tongue representation is spatially distant from the region of artifact and thus easy to distinguish.

To visualize the entirety of the somatomotor cerebellar topography, maps with representations of the four body part movements were projected onto a flatmap view of the cerebellum (Fig. 2). In this view, the inversion of the two map representations is clear. Also note that the second map evolves over the medial-lateral axis, with anterior-medial (close to the vermis) tongue representation distant from the posterior-lateral foot representation.

**Fig. 2.**
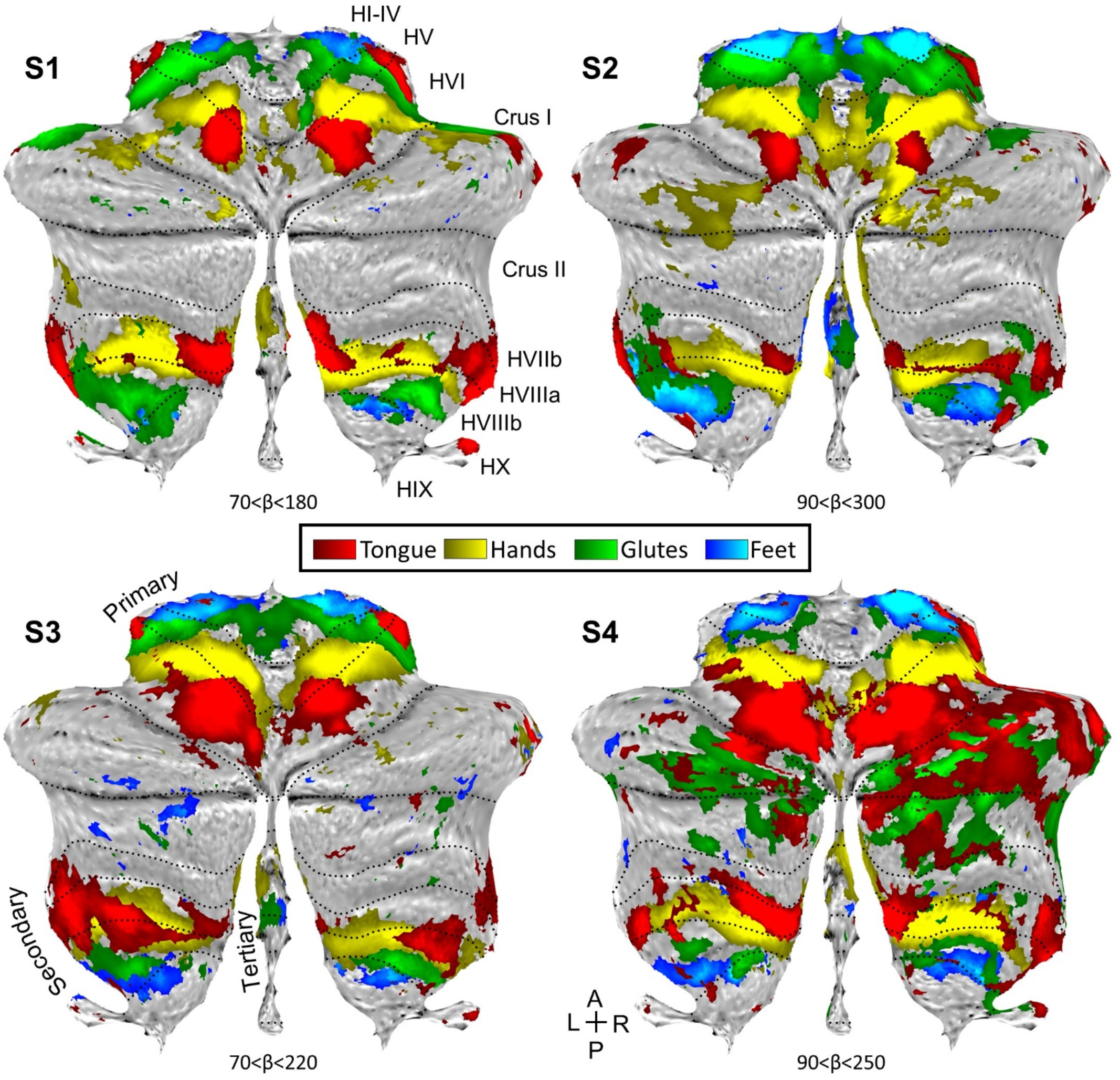
Somatomotor topography projected onto a flatmap of the cerebellum reveals details. Winnertake-all maps of active movements are displayed for 4 body parts along the anterior-posterior body axis projected onto flatmaps using the SUIT toolbox: tongue (red), right and left hands (yellow), glutes (green) and right and left feet (blue). Each panel displays a separate participant (S1-S4). In each participant, the primary and secondary somatomotor maps are apparent in the anterior and posterior lobes of each hemisphere corresponding to the anterior-posterior body axis. Also note the medial representations within the vermis (glutes and feet in S2 and S3, and hands in all four individuals). These may be hints for the hypothesized tertiary body representation in the cerebellum. Glutes and tongue responses in the Crus I / II of S4 are likely due to motion artifacts that are more common during these movements. Dotted lines indicate approximate lobule boundaries with lobules labelled for S1. L: Left; R: Right; A: Anterior; P: Posterior.

**Fig. 3.**
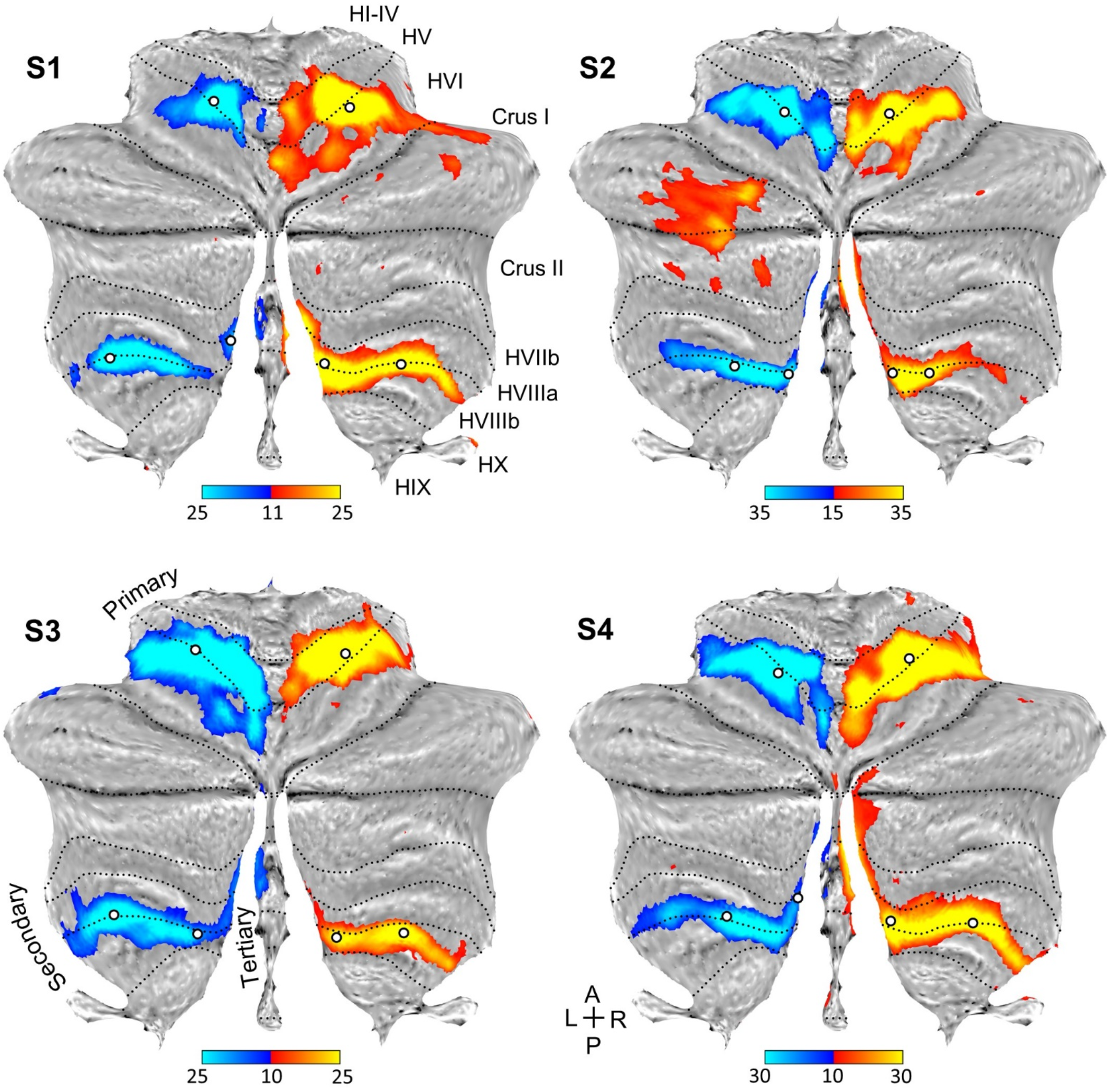
Direct contrast of left and right hand movements isolates representations of the body part. Contrast maps of right (red) versus left (blue) hand movements are projected onto flatmaps. Each panel displays a separate participant (S1-S4). In each participant, the hand representation is robust in both the anterior and posterior lobes. On the vermis, medial to the second representation, there is a bilateral hand representation in all participants. This extension of the hand representation may be a component of the tertiary representation, but it is ambiguous because there is no discontinuity. The white circles display the positions of seed regions that were used for analyses displayed in Fig. 5 (see text). Dotted lines indicate approximate lobule boundaries with lobules labelled for S1. L: Left; R: Right; A: Anterior; P: Posterior. Color bars indicate beta values (?)

Fig. 2 also shows provisional evidence for a third map within and near to the vermis. The hand representation in all four participants extends into the region of the vermis, as does the foot and glutes representations in S2 and S3. This feature is consistent with the hypothesized third (tertiary) somatomotor map of the cerebellum. However, idiosyncrasies in the data, variability across participants and the adjacency to the secondary representation make it uncertain. To further explore the possibility of a third map, we focused specifically on hand and foot movements where direct contrasts between the right and left movements were possible.

### The hand representation in the cerebellum is consistent with, but does not differentiate, a third somatomotor map

Contrasting right and left hand movements yielded robust responses in the cerebellum for all participants (Fig. 3). The primary and secondary representations were clear and distinct. In addition, all four individuals possessed bilateral hand representations on the edges of the vermis, medially and anteriorly to the second representation, potentially a component of the hypothesized third map. Specifically, in S1 and S2 there were two distinct representations for the second and the candidate third map. In S3 and S4 the two representations appeared as one continuous cluster. Consequently, it is still ambiguous as to whether the medial representation is a distinct third representation or simply a continuation of the second representation. It seems that even if there is a third map, the hand representation is not positioned to provide clear evidence for dissociation. Therefore, we repeated the same analyses focused on the foot representation.

### The foot representation in the cerebellum differentiates the third somatomotor map

Contrasting right and left foot movements yielded robust responses that included a representation in the vermis medial to the second representation in multiple participants (Fig. 4). Unlike the hand representation (Fig. 3), where the posterior lobe representation was continuous, the two representations associated with movement of the foot were spatially distinct (Fig. 4). One large representation was centered in lobule HVIIIb where the established second map’s upright topography begins. The smaller (newly) detected foot representations were medial and discontinuous with the known second somatomotor map. The presence of an independent foot representation in the vermis of the posterior lobe directly supports a third somatomotor map.

**Fig. 4.**
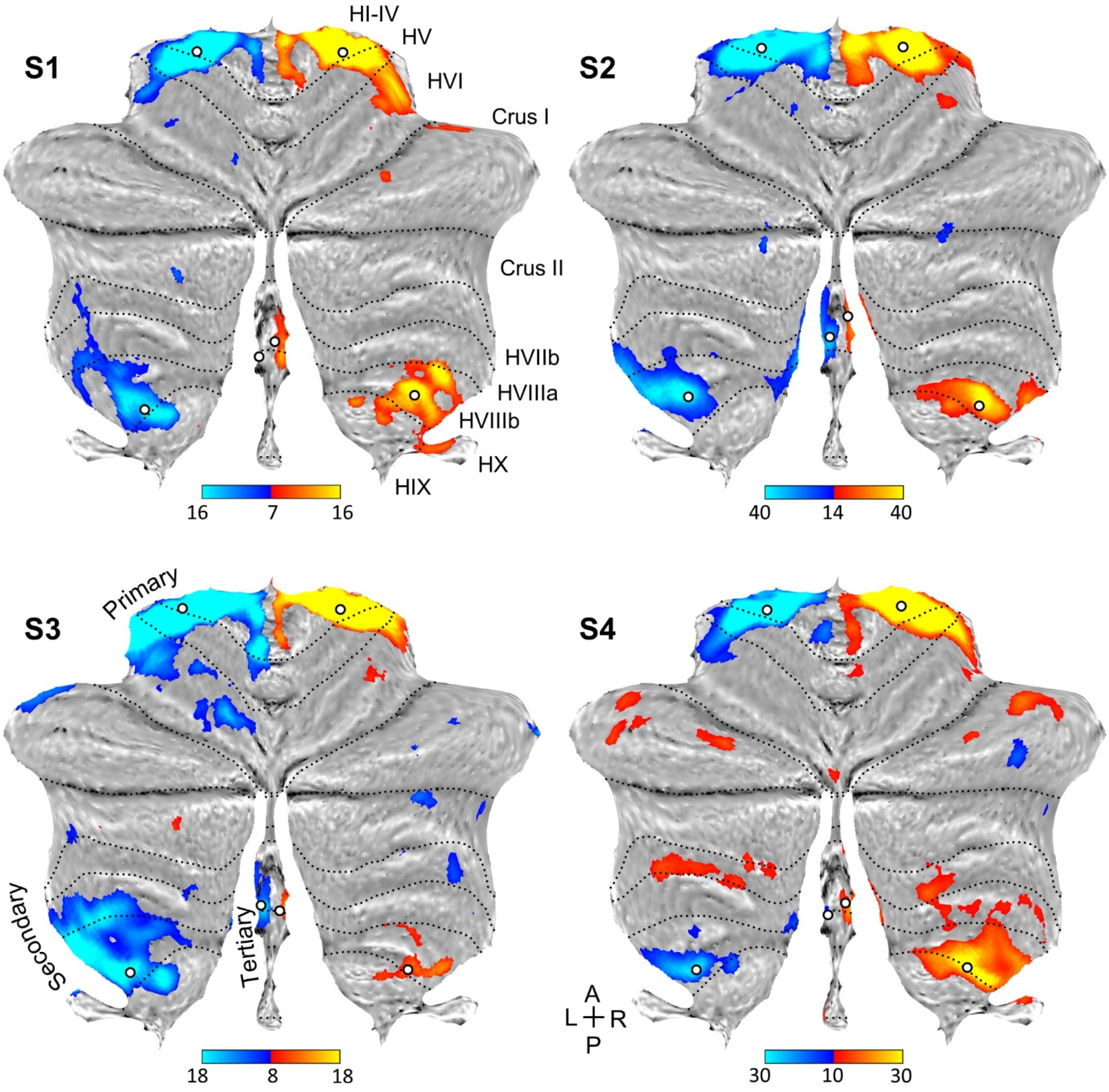
Direct contrast of left and right foot movements isolates three spatially discontinuous representations of the body part. Contrast maps of right (red) versus left (blue) foot movements are projected onto flatmaps. Each panel displays a separate participant (S1-S4). In each participant, the foot representation is detected in the anterior lobe and also the posterior lobe near to where the beginning of the secondary map is localized (laterally in lobule HVIIIb). Within the vermis, there is a distinct representation of the foot in each participant that is particularly clear in S2 and S3. This spatially discontinuous representation is evidence for a third somatomotor map. The white circles display the positions of seed regions that were used for analyses displayed in Fig. 6 (see text). Dotted lines indicate approximate lobule boundaries with lobules labelled for S1. L: Left; R: Right; A: Anterior; P: Posterior.

### The third cerebellar somatomotor map shows anatomically-specific coupling to cerebral motor cortex

Another way to refute or support the presence of independent cerebellar maps is to examine the specificity of cerebellar functional connectivity to the cerebral cortex. The positioning of the second and third somatomotor maps is such that (1) the spatially separate cerebellar foot representations should both couple to the same midline cerebral zone (see Fig. 1), and (2) the hand representations, which sit between the two foot representations in the cerebellum, should couple to a distinct cerebral zone near the hand knob of the precentral gyrus (see Fig. 1).

To explore these hypotheses, we first examined the specificity of the hand representations by applying seed-based functional connectivity analysis to the cerebellum and examining the correlation pattern in the cerebral cortex. Three sets of right and left seed regions were defined in the primary, secondary and candidate tertiary representations (white dots, Fig. 3). Given that the obtained posterior lobe hand representation does not possess a discontinuity to separate the second from the third maps, the seed regions were estimated to be at or near the transitions to the foot representations, but still within the hypothesized hand representation.

In all individuals, functional connectivity for all three seed region pairs revealed spatially-specific cerebral regions at or near the hand region of M1 (Fig. 5). Right cerebellar seed regions correlated with the left cerebral hand region and left seed regions with the right. The results for the tertiary seed regions were found to be the weaker but possessed the same, spatially-convergent pattern.

**Fig. 5.**
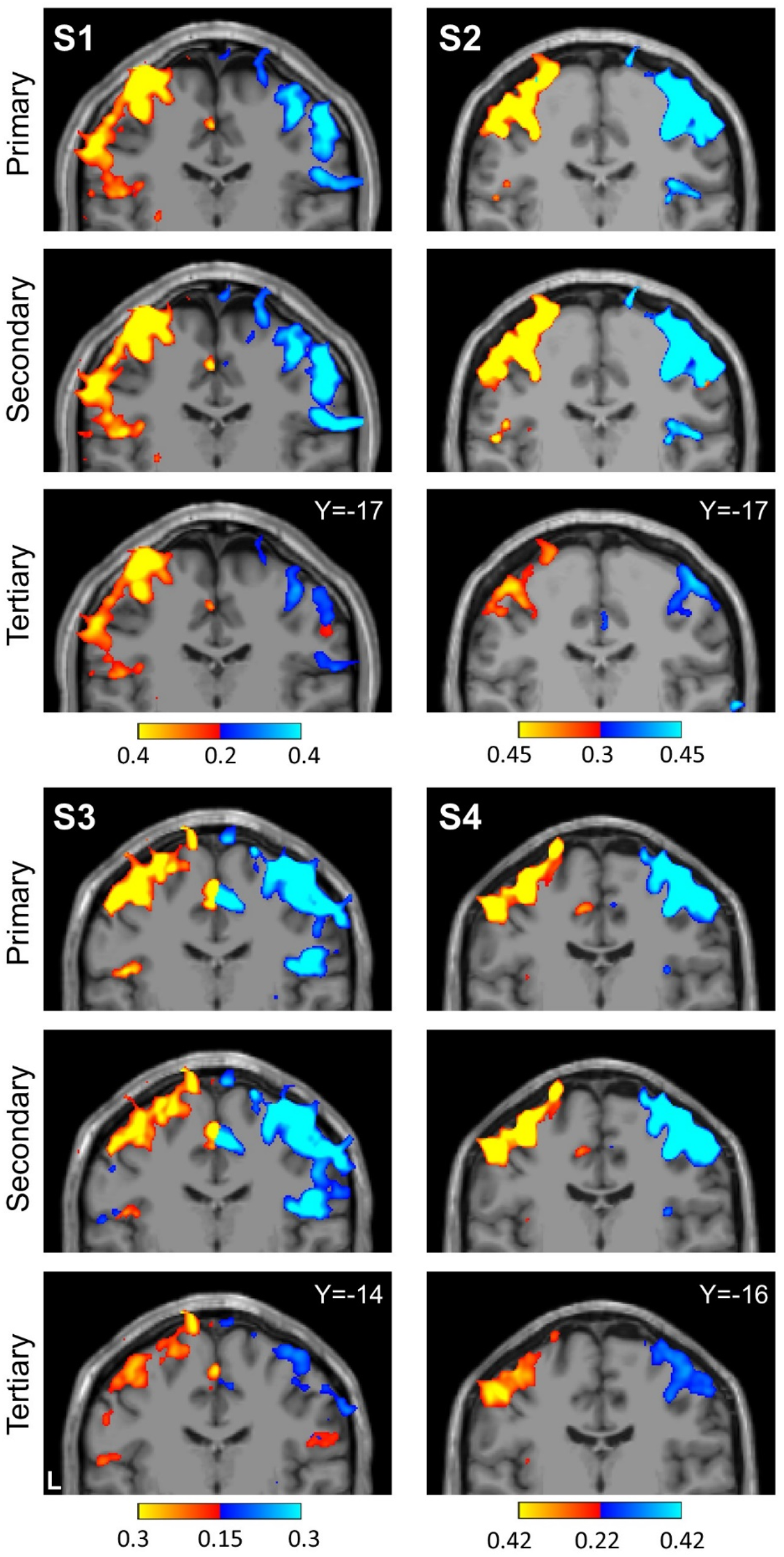
Seed based functional connectivity of cerebellar hand representations reveals the contralateral hand region in M1. Coronal sections display functional connectivity patterns for hand seed regions in the right (red) and left (blue) cerebellum. In each column of three panels, an individual participant’s data are shown for separate sets of right versus left hand region contrasts that are independently seeded in the three cerebellar representations. The locations of the seed regions are illustrated by white circles in Fig. 3. In each individual, functional connectivity resulting from the estimated primary, secondary and candidate tertiary cerebellar representations reveals M1’s contralateral hand region, demonstrating specificity. Note how, despite differences in correlation strength, the pattern revealed by the tertiary representation’s seed regions recapitulates the same pattern as the seed regions placed in the primary and secondary representations. Coordinates indicate the section level in the space of the MNI152 atlas. The color bars indicate correlation strength [z(r)]. L indicates left.

Applying a parallel analysis to the foot representations revealed a distinct pattern (Fig. 6). Unlike the hand representations, the seed region pairs for the second representation and candidate third representation of the foot are spatially discontinuous. Functional connectivity for all three seed region pairs revealed cerebral regions at or near the foot region of M1 that were distinct from the hand region. Of note, the hypothesized third foot cerebellar representation when seeded was, in isolation, able to recapitulate the selective cerebral motor pattern. This is important evidence because the zone of the cerebellum between the second and third maps is coupled to the cerebral hand region. The second and third foot representations are thus functionally selective and robustly dissociated from the cerebellar zone between them.

**Fig. 6.**
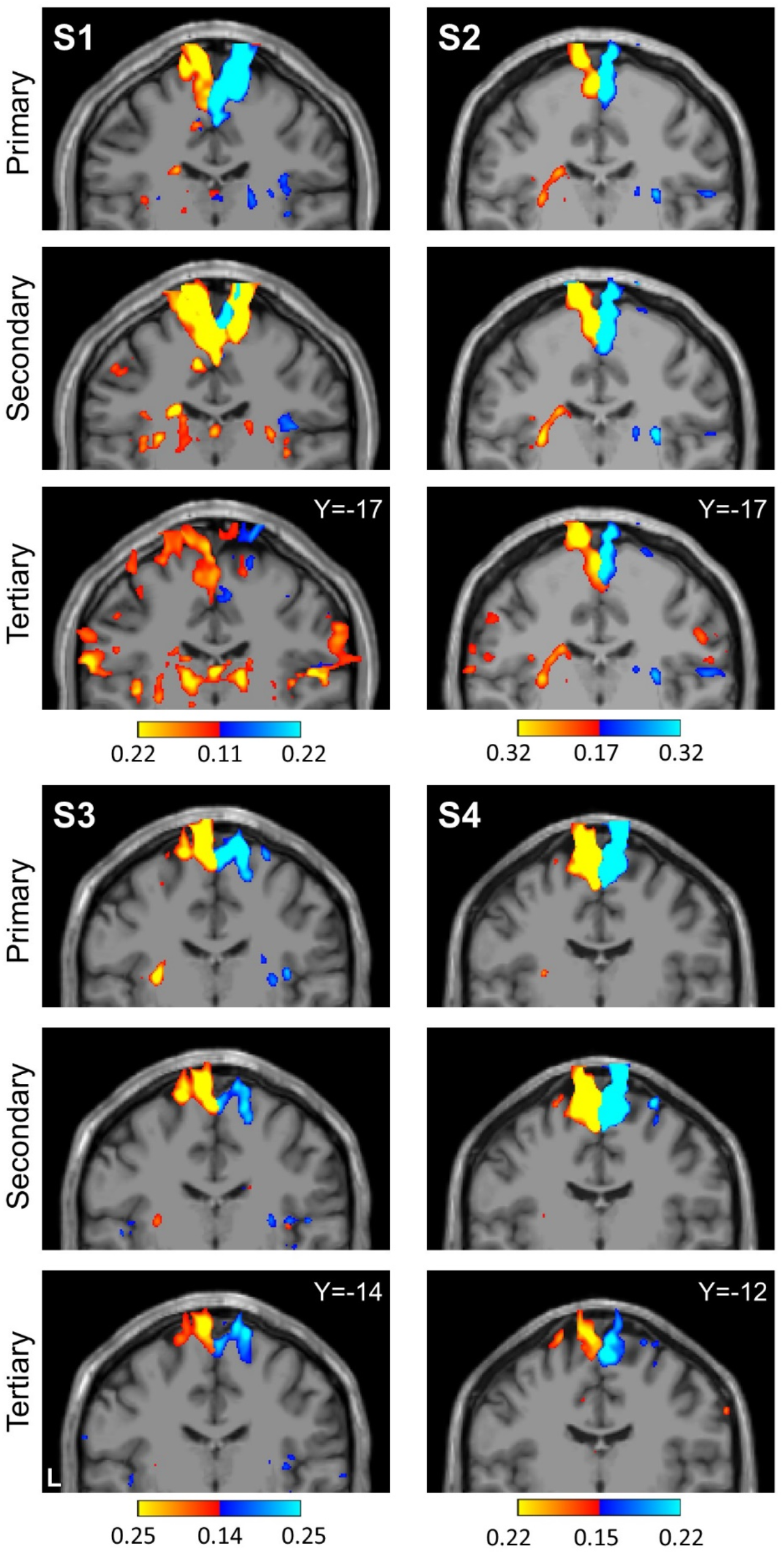
Seed based functional connectivity of cerebellar foot representations reveals the contralateral foot region in M1, including for the spatially discontinuous tertiary representation. Coronal sections display functional connectivity patterns for foot seed regions in the right (red) and left (blue) cerebellum. In each column of three panels, an individual participant’s data are shown for separate sets of right versus left foot region contrasts that are independently seeded in the three cerebellar representations. The locations of the seed regions are illustrated by white circles in Fig. 4. In each individual, functional connectivity resulting from the estimated primary, secondary and candidate tertiary cerebellar representations reveals M1’s contralateral foot region, demonstrating specificity. Note the pattern revealed by the tertiary representation’s seed regions recapitulates the same pattern as the seed regions placed in the primary and secondary representations despite being spatially discontinuous. Coordinates indicate the section level in the space of the MNI152 atlas. The color bars indicate correlation strength [Z(r)]. L indicates left.

To appreciate the discontinuity of the second and third somatomotor maps, the locations of the seed regions used in the above analyses were plotted on each individual’s structural volume image of their cerebellum. Seed regions for the second and third foot representations are on either side of the seed regions for the hand representation (Fig. 7). This observation alleviates concerns that the discontinuity could be an artifactual byproduct of the surface projection.

**Fig. 7.**
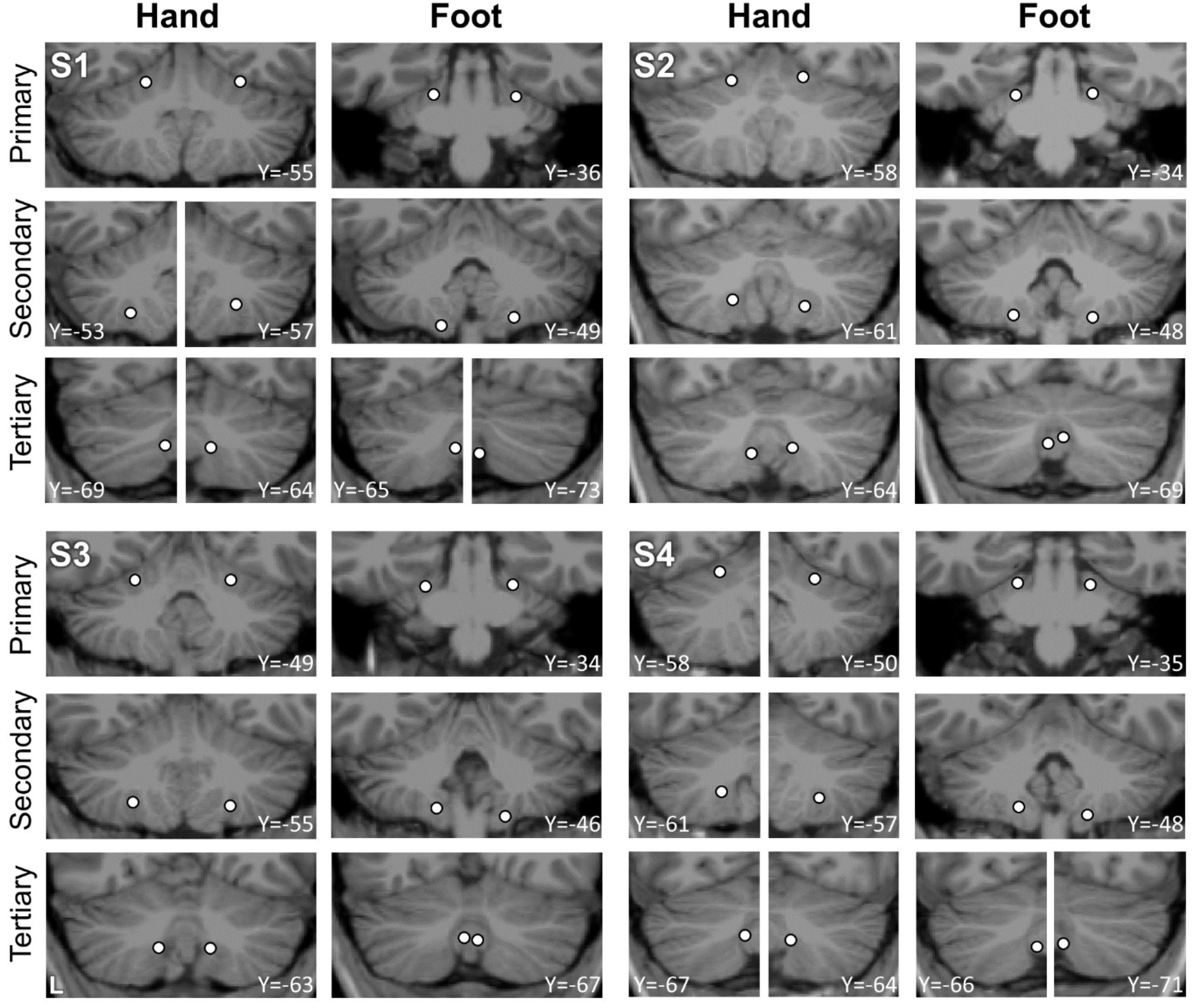
Visualization of seed region locations in the volume. Seed region locations of the right and left primary, secondary and tertiary representations (white circles) are plotted on coronal sections of the T1 structural image for each participant. Sections presented are of the seed region’s center or within 1mm of the center. For each individual, hand and foot seed region locations are presented. Note how the tertiary foot location is medial to the hand representation in each participant. Coordinates indicate the section level in the space of the MNI152 atlas. L indicates left.

**Fig. 8.**
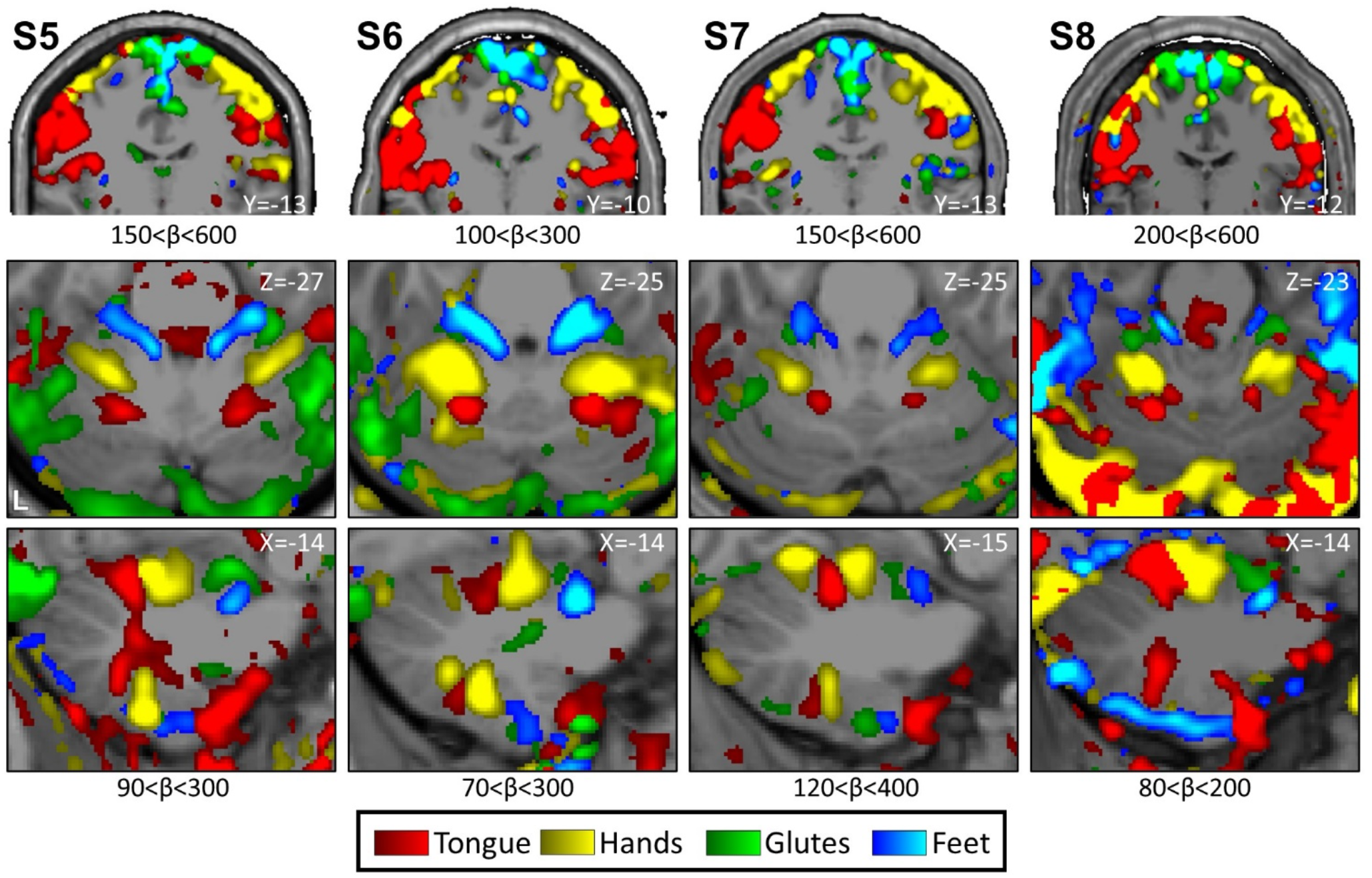
Replication of somatomotor topography in the cerebral cortex and cerebellum. Winner-take-all maps of active movements are displayed for 4 body parts along the anterior-posterior body axis: tongue (red), right and left hands (yellow), glutes (green) and right and left feet (blue). Each column displays a separate participant from the Replication sample (S5-S8). Analysis and plotting of these participants occurred after the results of the Discovery cohort were finalized. Beta values are thresholded to best capture topography, separately for the cerebral cortex and cerebellum. In each participant a clear body topography is evident by the order-tongue(T)-hand(H)-glutes(G)-foot(F) in the primary motor cortex, M1 (lateral to medial, top row), and cerebellar anterior lobe (middle and bottom rows). The primary cerebellar representation (T1-H1-G1-F1) is inverted to the secondary representation as labelled in S2 in Fig. 1 (T2-H2-G2-F2). S6 and S7 show a full topography also in the cerebellar posterior lobe (bottom row) while a partial topography is observed in S5 and only the tongue representation in S8. Coordinates indicate the section level in the space of the MNI152 atlas. The color bars represent beta values. L indicates left.

### Prospective replication of the third cerebellar somatomotor map

The findings above reveal strong evidence for a third somatomotor map in the cerebellum. Given the importance of this observation and also that we have previously failed to find clear evidence for a third map^1^, we sought to replicate all of the above observations in an independent set of new participants. Critically, none of the Replication sample data (S5 to S8) were processed until all of the analyses on the initial Discovery sample analyses (S1 to S4) were completed and plotted.

In the Replication participants, we again found evidence for full body topography in M1 (Fig. 8, top) and in the cerebellar anterior lobe of all four individuals with the expected order of the tongue-hand-glutes-foot (Fig. 8, top and middle rows, Fig.9). The second cerebellar map in the posterior lobe was also found with its full topography in S6 and S7. S5 and S8 were missing clear representation of the glutes. In participants S5, S6 and S7, responses consistent with a third map were also evident with foot representations present within the vermis (Fig. 9).

**Fig. 9.**
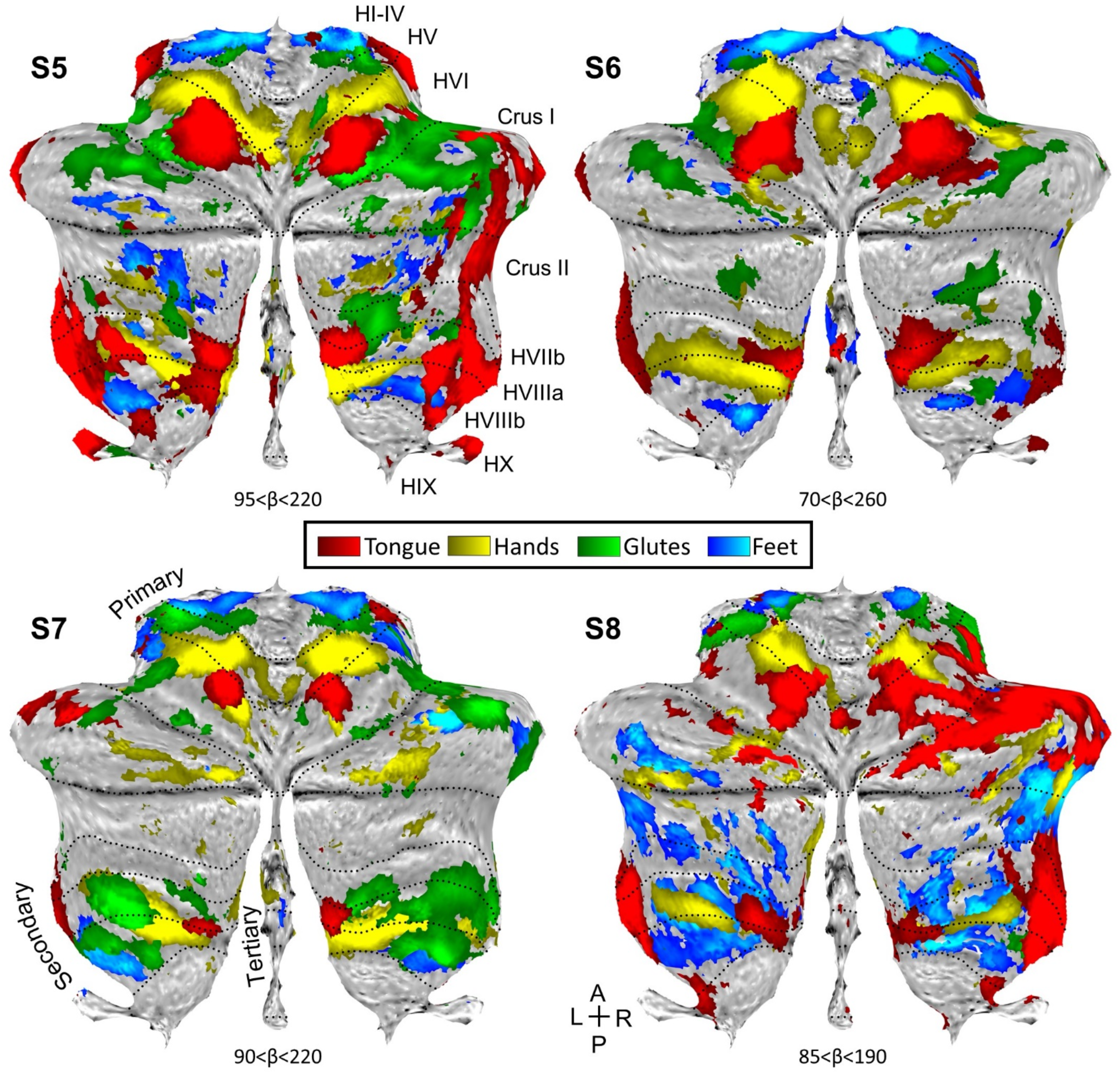
Replication of somatomotor topography projected onto a flatmap. Winner-take-all maps of active movements are displayed for 4 body parts along the anterior-posterior body axis projected onto flatmaps using the SUIT toolbox: tongue (red), right and left hands (yellow), glutes (green) and right and left feet (blue). Each panel displays a separate participant from the Replication sample (S5-S8). In participants S5-S7, the primary and secondary somatomotor maps are apparent in the anterior and posterior lobes of each hemisphere corresponding to the anterior-posterior body axis. The experiment largely failed in S8 (the primary map is apparent but the secondary map is ambiguous). Also note the medial representations within the vermis of S5-S7. These may be hints for the hypothesized tertiary body representation in the cerebellum. Dotted lines indicate approximate lobule boundaries with lobules labelled for S5. L: Left; R: Right; A: Anterior; P: Posterior.

Contrasting right and left hand movements again yielded robust responses in the cerebellum (Fig. 10). The maps of the hand representations were present in all participants. Repeating the same analysis for foot movements yielded evidence for the third somatomotor map with discontinuous representation of the foot near to the vermis (Fig. 11). S5 and S6 were particularly clean examples in the Replication sample, similar to the S2 and S3 in the prior Discovery sample. S8 showed a great deal of non-specific noise throughout the cerebellum and also lacked a clear foot representation within the vermis. Thus, across the two studies, there were multiple robust demonstrations of the third foot representation, but also the occasional ambiguous result and one clear failure (S8), consistent with prior studies of within-individual precision mapping that do not always achieve robust results in every individual (e.g., see Braga and Buckner 2017; DiNicola et al. 2020).

**Fig. 10.**
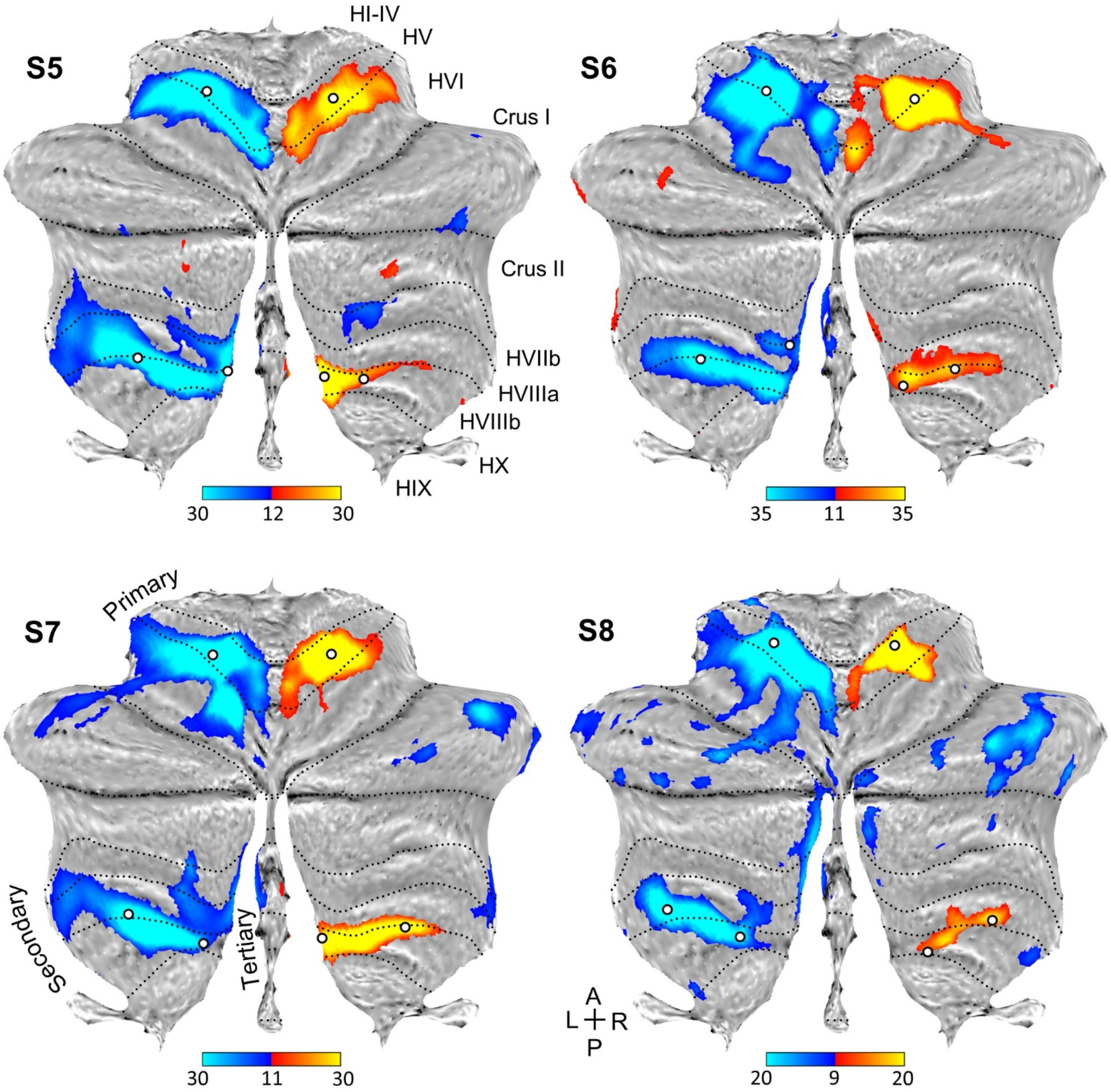
Replication of the hand representations. Contrast maps of right (red) versus left (blue) hand movements are projected onto flatmaps. Each panel displays a separate participant from the Replication sample (S5-S8). In each participant, the hand representation is robust in the anterior and the posterior lobes. The white circles display the positions of seed regions that were used for analyses displayed in Fig. 12 (see text). Dotted lines indicate approximate lobule boundaries with lobules labelled for S5. L: Left; R: Right; A: Anterior; P: Posterior.

**Fig. 11.**
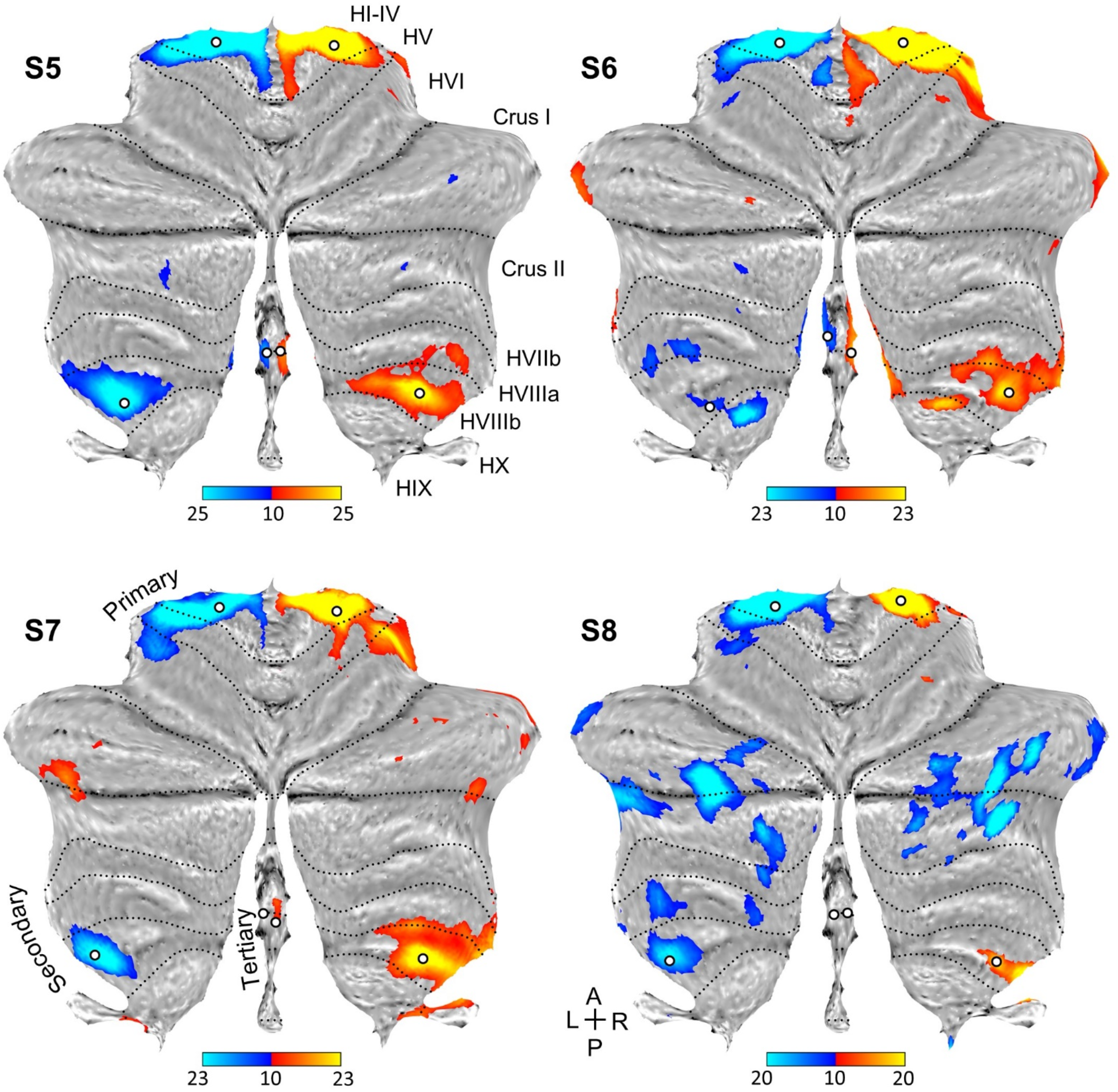
Replication of three spatially discontinuous foot representations. Contrast maps of right (red) versus left (blue) foot movements are projected onto flatmaps. Each panel displays a separate participant from the Replication sample (S5-S8). In three individuals, three separate foot representations can be identified - right and left primary, secondary and a tertiary representation in the vermis (only right in S7). In S8, a tertiary representation was not detected. The spatially discontinuous representation within the vermis replicates evidence for a third somatomotor map. The white circles display the positions of seed regions that were used for analyses displayed in Fig. 13 (see text). Dotted lines indicate approximate lobule boundaries with lobules labelled for S5. L: Left; R: Right; A: Anterior; P: Posterior.

Placing seed regions within each of the three cerebellar maps again revealed the expected anatomically-specific correlation patterns with cerebral motor zones (Figs. 12 and 13). High specificity was found in the three individuals for all three representations. S8’s connectivity pattern showed M1’s bilateral foot region from the primary and secondary seed regions (to lesser extent) but failed to demonstrate evidence for a tertiary representation. The locations of the seed regions are displayed in the volume (Fig. 14).

**Fig. 12.**
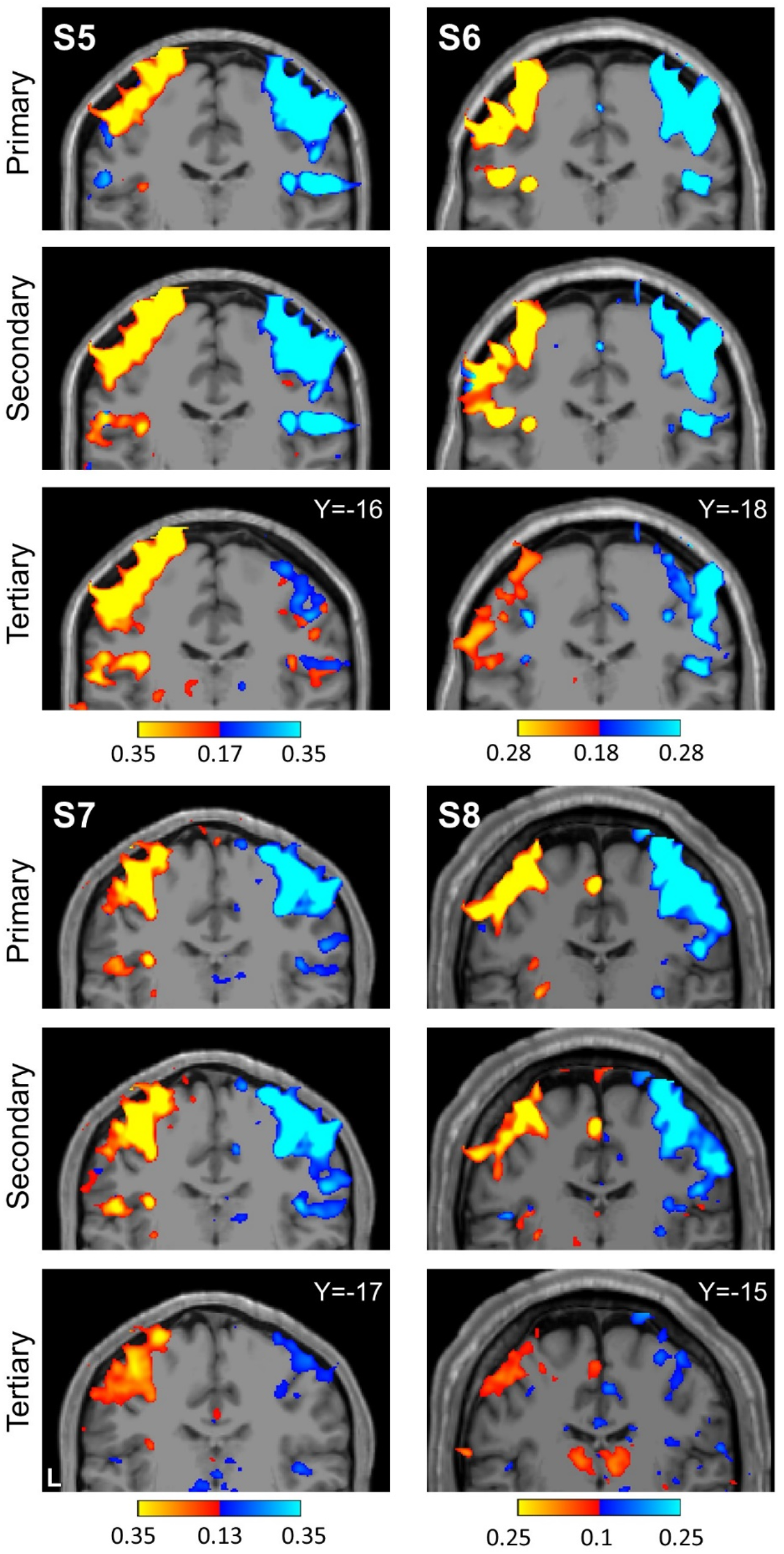
Replication of seed based functional connectivity from cerebellar hand representations. Coronal sections display functional connectivity patterns for hand seed regions in the right (red) and left (blue) cerebellum. In each column of three panels, an individual participant’s data from the Replication sample are shown for separate sets of right versus left hand region contrasts that are independently seeded in the three cerebellar representations. The locations of the seed regions are illustrated by white circles in Fig. 10. Functional connectivity resulting from the estimated primary, secondary and candidate tertiary cerebellar representations reveal M1’s contralateral hand region in all participants, demonstrating specificity. Note how, despite differences in correlation strength, the pattern revealed by the tertiary representation’s seed regions again recapitulate largely the same pattern as the seed regions placed in the primary and secondary representations. Coordinates indicate the section level in the space of the MNI152 atlas. The color bars indicate correlation strength [z(r)]. L indicates left.

**Fig. 13.**
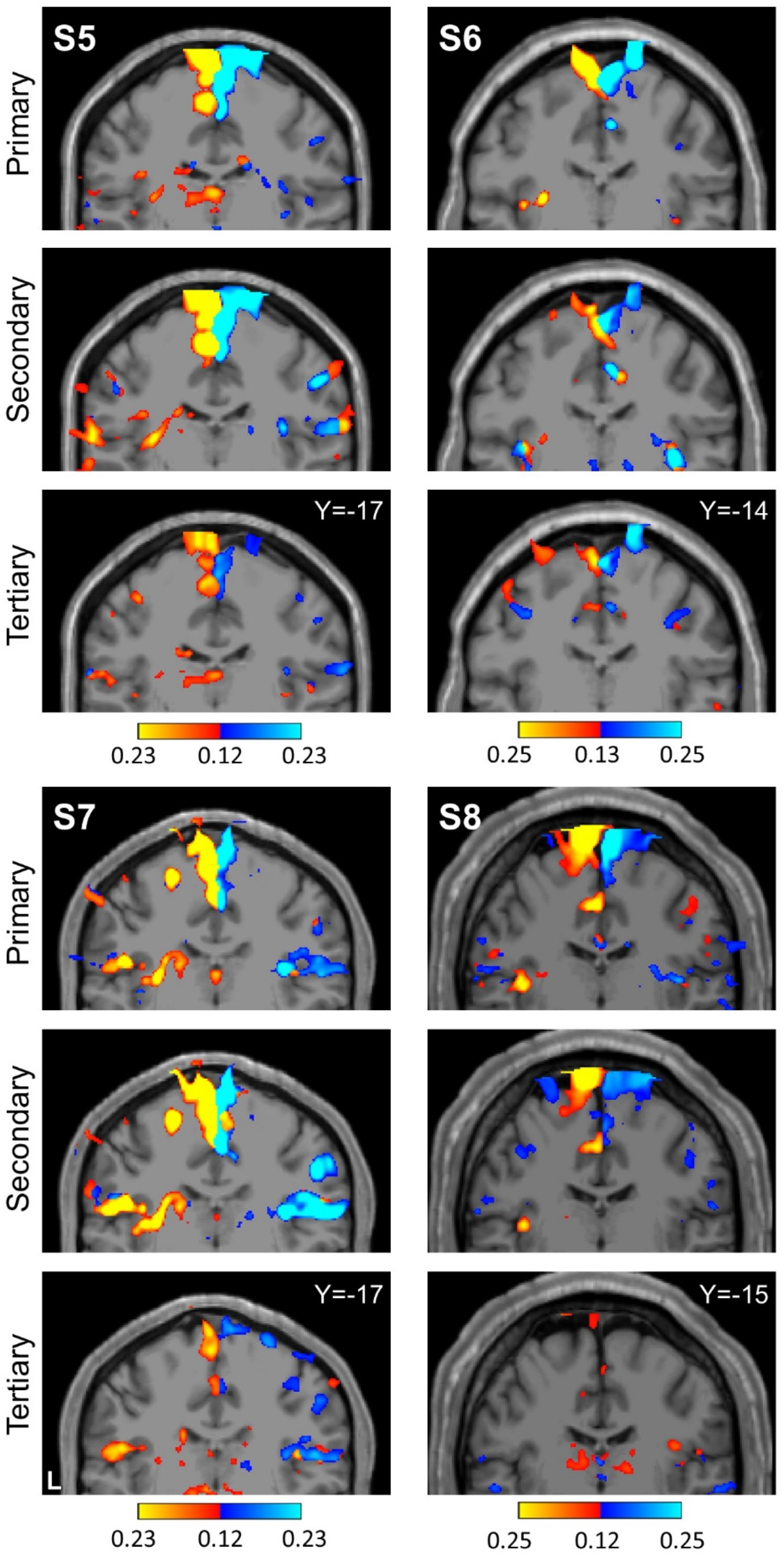
Replication of seed based functional connectivity from cerebellar foot representations. Coronal sections display functional connectivity patterns for foot seed regions in the right (red) and left (blue) cerebellum. In each column of three panels, an individual participant’s data from the Replication sample are shown for separate sets of right versus left foot region contrasts that are independently seeded in the three cerebellar representations. The locations of the seed regions are illustrated by white circles in Fig. 11. In three individuals (S5-S7), functional connectivity of seed regions in the primary, secondary, and candidate tertiary cerebellar representations reveal M1’s contralateral foot region (with S5 and S6 clearer than S7). The pattern is absent in S8. Coordinates indicate the section level in the space of the MNI152 atlas. The color bars indicate correlation strength [z(r)]. L indicates left.

**Fig. 14.**
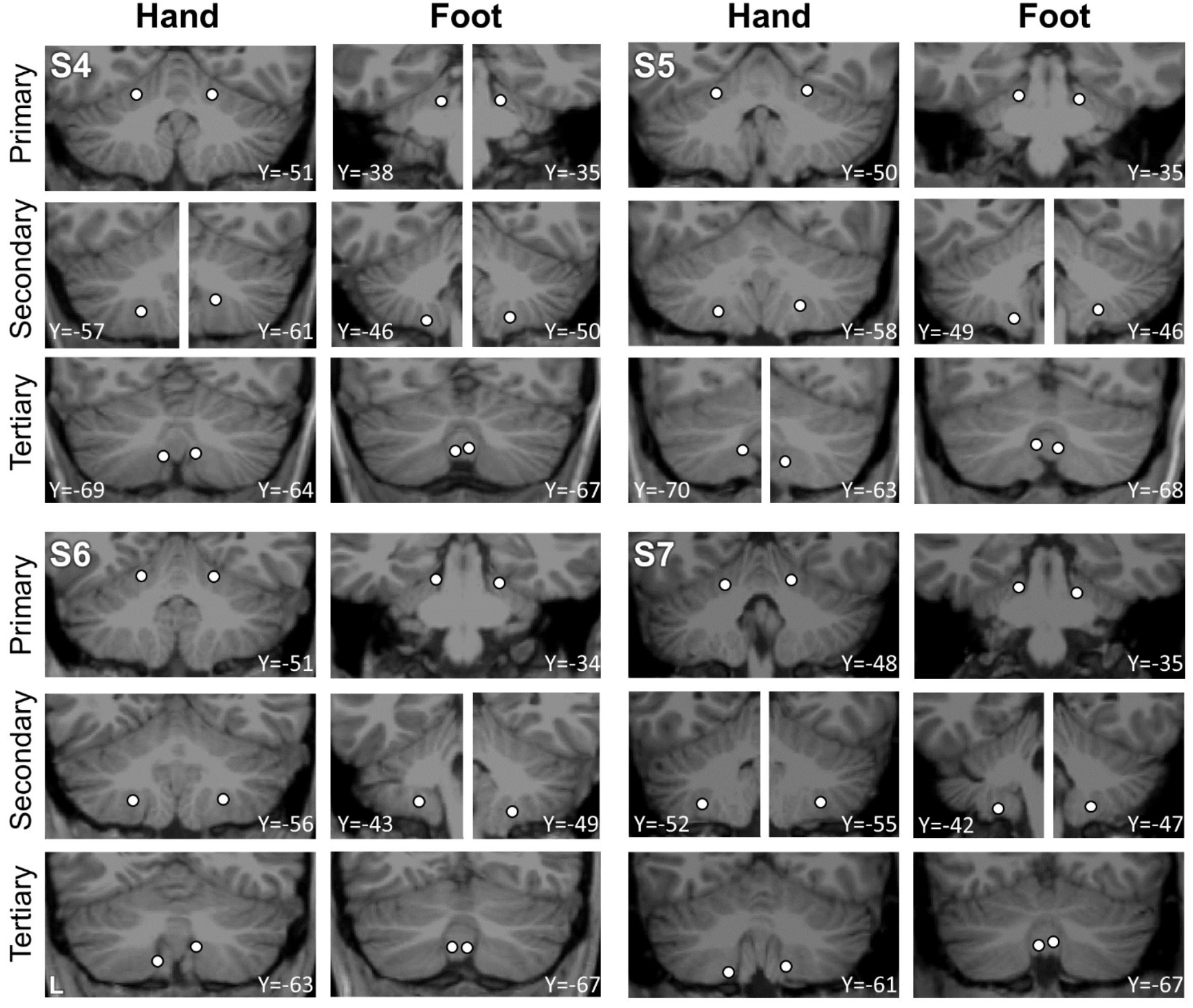
Visualization of seed region locations in the volume for S5 to S8. Seed region locations of the right and left primary, secondary and tertiary representations (white circles) are plotted on coronal sections of the T1 structural image for each participant in the Replication sample. Sections presented are of the seed region’s center or within 1mm of the center. For each individual, hand and foot seed region locations are presented. Note how the tertiary foot location is medial to the hand representation in each participant. Coordinates indicate the section level in the space of the MNI152 atlas. L indicates left.

### The three somatomotor maps are evident when examined at the group level

While our studies were designed with the goal of preserving idiosyncratic anatomical details within the individual, the results suggested that there may be sufficient positional stability of the three somatomotor maps to examine the data at the group level. Fig. 15 (top) shows positions of hand and foot primary, secondary and tertiary seed regions of all 8 individuals overlayed on a flatmap. Three foot representations separated by the hand are clearly present suggesting that averaging data across participants can be effective.

**Fig. 15.**
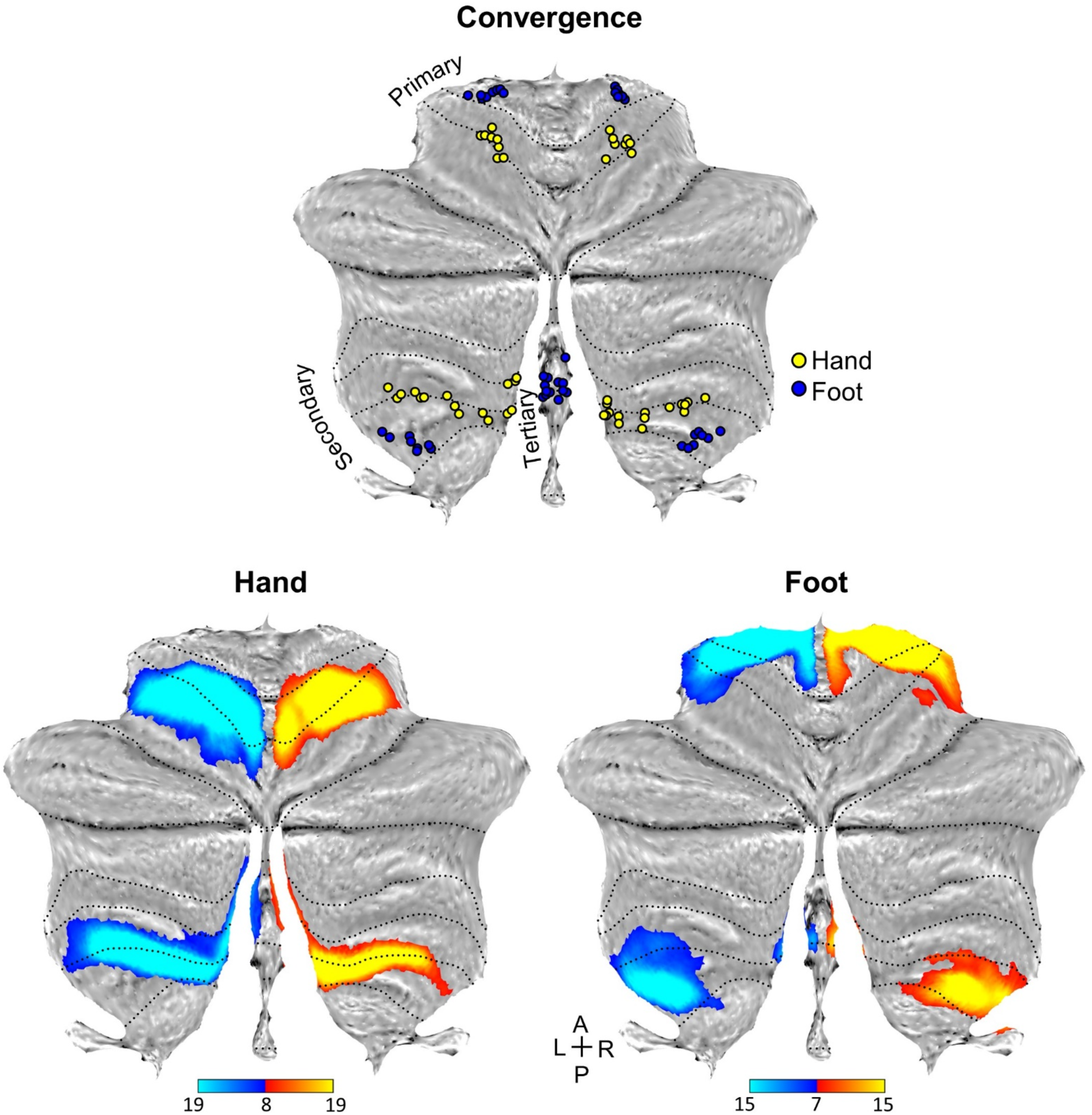
The three somatomotor representations are detected in the group-averaged flatmap of the cerebellum. (Top) Hand and foot primary, secondary and tertiary seed regions from all 8 participants are projected on a flatmap. Note the three separated foot representtions with the hand representation between the second and candidate third map. (Bottom left) A contrast map of right (red) versus left (blue) hand movements, averaged across participants, is projected on a cerebellar flatmap. The map is the mean beta map of all 8 participants including 192 separate runs collected during active movements. Dotted lines indicate lobular boundaries as labelled in Fig. 2. Note the robust separation of the primary and secondary representation of the hand, with the secondary representation extending to the vermis. (Bottom right) A parallel contrast map of right (red) and left (blue) foot movements is displayed. Note that there are three separate representations of the foot, with a tertiary representation falling within the vermis. The tertiary representation is spatially discontinuous with the secondary representation. The color bar represents the mean beta values. L: Left; R: Right; A: Anterior; P: Posterior.

Fig. 15 (bottom) shows the left versus right contrasts of the hand and foot movements after averaging the beta values across all 8 individuals (combining all data from the Discovery and Replication samples). Critically, the small discontinuous responses linked to the right and left foot movements are evident within the vermis as would be expected from the analyses of the individuals. Once the location is known, it is now possible to see evidence in the group data. Fig. 16 illustrates the location of the third representation of the foot in the group-averaged volume (marked by asterisk). The positioning and small size of the response also suggest why the third map representation is challenging to detect (and notice) as separate. We will return to this point in the discussion where the existing literature was reexamined for prior overlooked evidence for the third map.

**Fig. 16.**
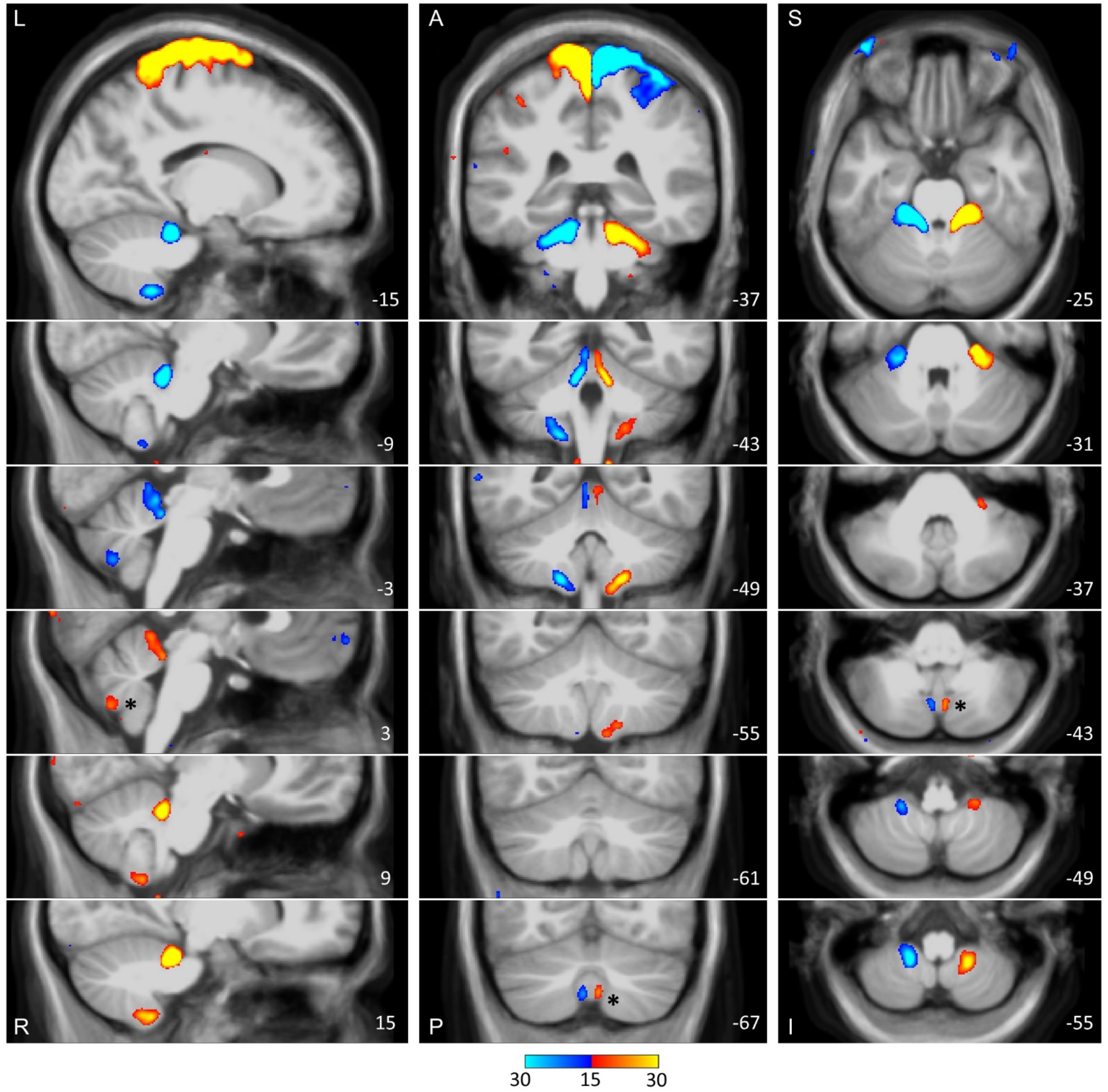
The three somatomotor representations of the foot are anatomically distinct in the group-averaged volume of the cerebellum. The three columns display sections in sagittal (Left), Coronal (Middle), and transverse (Right) orientation. In each orientation, the third foot representation is detected and anatomically separated from the second foot representation. The right third representation is marked by an asterisk in each view and is smaller in size as compared to the other two representations. The anatomical backdrop is the average volume from the 8 participants contributing the functional data. In the middle and right columns, left is displayed on the left. Coordinates at the bottom right of each panel indicate the section level in the space of the MNI152 atlas. The color bar represents the mean beta values. L: Left; R: Right; A: Anterior; P: Posterior; S: Superior; I: Inferior.

## Discussion

Our data provide evidence for a third somatomotor map near to the vermis of the human cerebellum. Unlike the two well-established somatomotor maps that can be distinguished as separate by exploring the hand, foot or other body parts bilaterally, the linchpin for detecting the separation between the second and third maps was examining the multiple representations of the foot. When left and right foot movements were contrasted, a response was detected in the cerebellar vermis that was spatially discontinuous from the known second map. Moreover, the region of the posterior cerebellum between the two foot representations was associated selectively with the hand, providing evidence that the two foot representations are parts of spatially- and functionally-distinct maps. All findings were detected in multiple independent participants and replicated in a second, prospectively analyzed cohort. We discuss these observations in the context of what they reveal about the broad topography of the cerebellum as well as implications for the specific organization of the vermis.

### Evidence for the elusive third cerebellar somatomotor map

While two somatomotor maps within the cerebellum have long been established (Adrian 1943; Snider and Stowell 1944; Snider and Eldred 1952; for review see Manni and Petrosini 2004), it has been difficult to find clear evidence for a third map^1^. Guell and colleagues hypothesized that a third somatomotor map may not exist, suggesting an organizational distinction between how motor zones of the cerebellum are represented in contrast to nonmotor zones (Guell et al. 2018a, 2018b). By their “double motor / triple nonmotor representation hypothesis”, the most posterior zones of the cerebellum are occupied only by regions associated with higher-order cognitive and affective functions. The present results suggest that three distinct somatomotor maps may be present in the cerebellum, with the third falling at or near the third representation of nonmotor networks.

There are multiple possible reasons why the third map representation has been challenging to identify. First, it is small and buried in a difficult to access and hard to image region of the cerebellum. Second, studies using human neuroimaging have tended to focus on hand and finger movements and less on direct contrasts of foot movements. For example, the recent work by King and colleagues that produced detailed, comprehensive maps of cerebellar functional zones in both groups and individuals included only finger movement paradigms and did not include foot movements (King et al. 2019). We suspect if detailed analyses of foot movements had been more common in the field, consensus about the representation may have already emerged.

A third challenge deserving deeper discussion is that task-free functional connectivity estimates of cerebellar motor representations have been surprisingly equivocal. As one example from our own recent work where we explored two individuals each scanned across 31 sessions, the motor organization was the weakest aspect of the data. Even after revisiting that data based on the present findings, we cannot clearly establish the second map by contrasting left and right motor regions linked to the feet, despite there being little debate about the existence of the second map. The third map, being smaller than the second, is not detected. As another example, in our studies at high field (e.g., Braga et al. 2019) we were not able to detect robust functional connectivity between cerebral motor zones and the cerebellum in the face of clear functional connectivity from higher-order networks including the default network. In the recent work of Marek et al. (2018) that examined within-individual estimates of cerebellar organization from extensive functional connectivity data, the second map is variably absent. This can be seen in individuals but also in their averaged flatmap representation (see Marek et al. 2018, their Fig. S2). While there is a clear inverted topography in the anterior lobe, posterior lobe cerebellar regions within the vicinity of the established second somatomotor map are assigned to higher-order cognitive networks including the default network. We suspect functional connectivity between the cerebral cortex and the cerebellum, as examined in our multiple earlier studies and also in these other studies in the literature, is yielding an incomplete estimate of somatomotor organization. We do not fully understand the origins of this limitation but have been perplexed previously on multiple occasions^1^.

In this light it is interesting to revisit the results reported in Guell et al (2018a) because their study is one of the few to examine cerebellar organization using both task-free functional connectivity and also task-evoked active motor movements. Using data from the Human Connectome Project (Van Essen et al. 2013), the study is well-powered, including data from 787 participants with the ability to directly contrast left and right active foot movements. What is notable is that the group-averaged task-evoked estimates in Guell et al. (2018a, their Fig. 4) are highly similar to our present group-averaged map (Fig. 15), with a spatially-discontinuous representation of the foot in the posterior vermis. The estimate of the foot representation from the task-free data does not show the same pattern as the task-evoked estimate. Thus, when examined closely, the present results converge with earlier studies that have examined task-evoked somatomotor topography.

### Contextualizing the third somatomotor map within the broader organization of the cerebellum

The cerebellum possesses multiple roughly homotopic representations of the full cerebrum (Buckner et al. 2011b; see also Buckner et al. 2013; Guell et al. 2018b; Xue et al. 2021). Two sets of somatomotor, cognitive and affective networks are found that begin in the anterior lobe and progress along the anterior-to-posterior axis. A primary representation spans from the anterior extent of the cerebellum to Crus I / II, and a secondary representation spans from Crus I / II to the posterior extent of the somatomotor representation^2^. Yet, a parsimonious account of cerebellar organization as multiple repeating cerebral representations encounters an inconsistency.

Evidence for a third partial representation of cerebral networks has been found posterior to the second somatomotor representation (Buckner et al. 2011; Guell et al. 2018a; King et al. 2019; Xue et al. 2021). The third representation includes cognitive and affective regions, but the expected somatomotor component that would complete the representation has been missing. The present finding of a third foot representation, localized to the posterior vermis (Figs. 15 and 16), is consistent with the possibility that the cerebellum possesses three separate representations of the full cerebrum each including a topographic representation of somatomotor cortex.

The location of the third map within the vermis is particularly intriguing because this region has been understudied in humans despite being a major focus or work in animal models (e.g., Peters and Monjan 1971; Optican and Robinson 1980; Yamada and Noda 1987; Takagi et al. 1998; Herzfeld et al. 2015). Early studies reported responses to auditory stimulation in the cerebellar vermis of cats and monkeys (Snider and Stowell 1944; Huang and Liu 1991). Recently, van Es and colleagues demonstrated that the human cerebellar vermis responds to visual stimuli (van Es et al. 2019). Detailed within-individual analysis confirmed this observation and further noted that the region is coupled to primary retinotopic visual cortex (Xue et al. 2021). The third foot somatomotor representation is spatially near to the visual and auditory representations within the vermis, suggesting that multiple sensory and motor maps may be proximal to one another in this small buried region of the cerebellum, and also near to representations of nonmotor cerebral networks, including those important to affective function (e.g., Schmahmann and Sherman 1998; Stoodley and Schmahmann 2009). This juxtaposition of sensory and motor maps between domains is not predicted by a simple model in which all aspects of cerebellar topography are understood within a continuous homotopic mapping of the cerebrum, as visual cortex is distant from somatosensory and motor cortices in the cerebrum. Further study of the organization and functional importance of the human vermis is thus of great interest.

### A gap remains

Our data provide direct evidence for three separate discontinuous representations of the foot. However, despite mapping four body parts extensively within each individual participant, our results failed to reveal the orientation of the third map. There are multiple possible reasons for this gap. First, the tertiary representation is likely smaller than the other two. It is possible that the spatial resolution used here is not sufficient for mapping the full topography, especially given the complex surface topology of the cerebellum (Sereno et al. 2020). Second, the tongue and the glutes are more susceptible to head motion, and therefore the corresponding representations are noisier than the ones for the hands and feet, which also allow direct subtraction of left and right movements. Future explorations with intensive within-individual sampling at higher resolution will be informative. In addition, exploring a somatosensory paradigm where lateralized stimulation (right versus left) can be administered repeatedly across multiple body parts along the anterior-posterior axis may provide a path forward.

### Conclusions

We provide reliable evidence that the human cerebellum possesses three spatially-distinct representations of the foot. The location of the third foot representation is in the vermis, consistent with a third somatomotor map continuous with the tertiary representations of cognitive and affective networks. The third map’s small size and location may have contributed to its evasiveness in past explorations, and suggests examination at high spatial resolution will be necessary to fully understand the third map’s spatial orientation and its relation to functionally-distinct zones of the vermis.

## Acknowledgments

K. Ntoh, E. Iannazzi and A. Hamadeh assisted with data acquisition, and S. Kaiser assisted with data preparation and upload for open release. We thank the Harvard Center for Brain Science neuroimaging core and FAS Division of Research Computing. We thank T. O’Keefe for assisting in optimization of data processing, R. Mair for MRI physics support, and J. Diedrichsen for providing the SUIT toolbox. The multiband EPI sequence was generously provided by the Center for Magnetic Resonance Research (CMRR) at the University of Minnesota.

## Footnotes

1 Within our own work we have previously looked for a third map using functional connectivity fMRI both at high-field, high-resolution 7T (Braga et al. 2019) and within intensively studied individuals at 3T (Xue et al. 2021). These efforts yielded equivocal results. Considering also Buckner et al. (2011), the present effort is our fourth attempt to find evidence for a third somatomotor map.

2 We heuristically term the somatomotor / body maps as primary, secondary, and tertiary. As discussed by Guell et al. (2018a), this terminology is a heuristic for convenience and should not be taken to imply a known relation between the representations such as exists in primary and secondary visual cortical areas where direct thalamic input goes to the earlier (primary) area in a hierarchy of anatomically-connected areas. The ordering of the somatomotor maps in the cerebellum reflects the order in which the representations were discovered, not a functionally meaningful hierarchy.

